# Sizing single trapped nanoparticles with interferometric scattering fluctuations

**DOI:** 10.64898/2025.12.23.696206

**Authors:** Abhijit A Lavania, William Carpenter

## Abstract

The sizes of nanoparticles significantly influence their properties but are challenging to quantify at the single-particle level. To enable label-free single-nanoparticle detection, interferometric scattering microscopy has emerged as a sensitive modality that exhibits high-precision axial information encoded in the interferometric scattering signal. Here, we demonstrate that single nanoparticles in an interferometric scattering anti-Brownian electrokinetic (ISABEL) trap can be hydrodynamically sized by interferometic scattering contrast fluctuations. The fluctuation timescale is characterized by time autocorrelation analysis and interpreted via a Brownian Dynamics model of the ISABEL trap to provide an estimate of the hydrodynamic diameter of each trapped particle. We verify this behavior for gold and polystyrene bead standards, and demonstrate performance on single carboxysomes, nanoscale carbon fixation compartments from autotrophic bacteria. The exquisite sensitivity of interferometric scattering amplitude fluctuations further equips researchers with a label-free optical platform for increasingly sophisticated nanoparticle analysis.

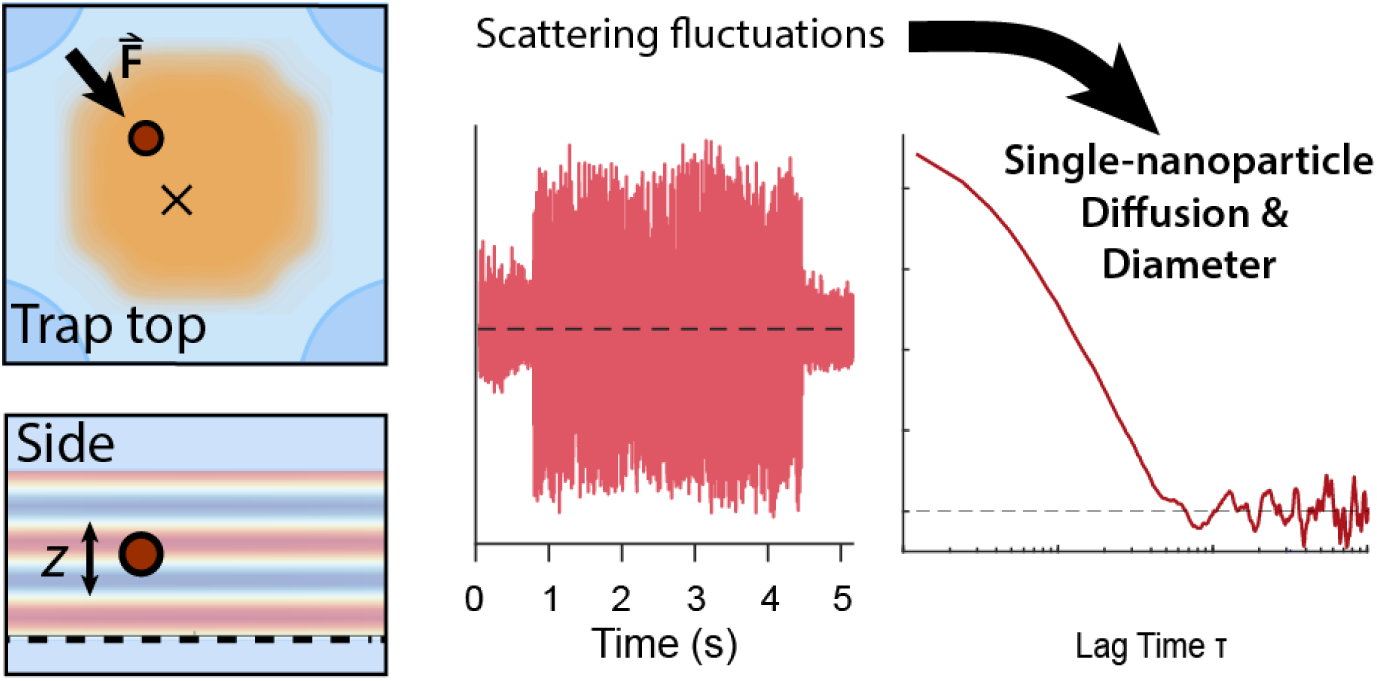

## 1. Introduction

Sophisticated nanoparticles are revolutionizing biochemical control at the nanoscale,^1–3^ where the sizes of these particles play a substantial role in their efficacy.^4^ In pharmaceuticals, nanoparticle size impacts *in vivo* distribution and circulation,^2^ drug release rate,^5^ and particle uptake even at the single-cell level.^6^ As sensors, the size-dependent surface plasmon resonances of metallic nanoparticles enable single-analyte detection and microbial activity in the biomedical,^1^ environmental monitoring,^7^ and agricultural sectors.^8^ With the increasing prevalence of multi-step nanoparticle synthesis and processing, size distribution can evolve at each step, leading to heterogeneity of performance.^9^ Because of this important design parameter, measuring nanoparticle diameter accurately and precisely is key to validation.^10^

Because nanoparticle diameter is a particularly challenging parameter to measure, distinct techniques quantification with varying strengths and limitations are in use. Optical microscopy is impractical since these nanoparticles are smaller than the optical diffraction limit ∼λ/2^11^ and super-resolution microscopy requires a demanding and slow acquisition.^12^ While the high spatial resolution of electron microscopy permits direct image analysis of each nanoparticle,^13^ the substantial effort and training needed for this technique presents considerable barriers to usage. Instead, solution-phase techniques like dynamic light scattering (DLS),^14^ fluorescence correlation spectroscopy (FCS),^15,16^ and nanoparticle tracking analysis (NTA)^17^ use the Stokes-Einstein equation to relate diffusional motions to effective hydrodynamic diameters *d*_H_. Despite relatively straightforward workflows, FCS and DLS produce ensemble-averaged values, which can misrepresent the distribution of sizes within a sample.^18^ Nanoparticle tracking analysis utilizes single-particle microscopy to directly observe the Brownian trajectories of individual nanoparticles, but short observation windows artificially broaden the retrieved nanoparticle size distribution.^10,19^

To address the above limitations in nanoscale measurement, interferometric scattering microscopy has recently demonstrated remarkable sizing performance.^20–22^ This is achieved by the sensitivity in the interference between a nanoparticle scatterer and a back-reflection from a nearby buffer-glass interface.^23–25^ In this scheme, the amplitude of the signal scales linearly with the volume of the nanoparticle^23,25^ for particles of uniform composition (versus squared-volume scaling in dark-field scattering and NTA^25^), drastically enhancing signal for particles below ∼30 nm in diameter.^23^ This label-free and non-destructive technique has been exploited for high-speed single-particle tracking,^26^ label-free cellular microscopy,^27,28^ and single-protein detection.^29^ Combined measurements of both Brownian trajectories and scattering contrast enable estimation of hydrodynamic diameter and effective refractive index of single biomolecular particles separately,^22^ enabling biomolecular composition analysis and particle discrimination in complex mixtures.^21,30^

Additionally, the phase difference between the scattered and reflected electric fields is a sensitive reporter of the particle’s height above the interface. For dielectric nanoparticles, a significant contribution to this phase is the optical path difference between the nanoparticle and the interface.^31,32^ As a particle diffuses axially, the signal fluctuates both in its sign and magnitude.^22^ Due to the phase redundancy in the axial direction, direct readout of the particle’s height is possible but non-trivial.^33^ Alternatively, time autocorrelation analysis is comparatively straightforward for quantifying a diffusional fluctuation timescale^14^ and is extensible to complex environments such as in live cells.^34^ Intensity time autocorrelation analysis is a familiar tool in single-particle science, additionally employed in single-molecule detection by fluorescence microscopy^16,35^ and cavity resonance fluctuations.^36,37^ Finally, time autocorrelation analysis forms the basis for differential dynamic microscopy,^38,39^ where fluctuations in optical signal from a microscopic scattering body reveal its underlying nanoscopic dynamics.

To extend the monitoring timescale of single nanoparticles and enable multiplexed spectroscopy, we recently developed the interferometric scattering anti-Brownian electrokinetic (ISABEL) trap.^40,41^ In this trap, we hold a single nanoparticle in solution for up to several minutes by applying active electrokinetic feedback to an individual nanoparticle’s position.^42^ The ISABEL trap emerged as a generalization of the fluorescence-based ABEL trap,^43,44^ demonstrating that we can apply this active trapping approach to single nanoparticles label-free.^40^ Importantly, the ISABEL trap enables a productive synergy between simultaneous scattering and fluorescence spectroscopic measurements to observe coupled biochemical and nanophysical properties in biomolecular nanoparticles.^45,46^ As a consequence of detecting particles with interferometric scattering, we observe large fluctuations in the sign and magnitude since the particle diffuses axially over ∼ 1μm.

Here, we show that we can estimate a single trapped particle’s axial diffusion coefficient and hydrodynamic diameter via time autocorrelation analysis. The observed contrast fluctuations deviate qualitatively from the expectation based on analytical models, so we developed a Brownian Dynamics simulation of the ISABEL trap to account for the experimental factors in the trap. By generating a library of modeled particles with varying diameters and scattering contrasts, we can compare an experimentally determined ACF to this library to estimate a single particle’s hydrodynamic diameter to within a few nm. We validate this approach with standard gold and polystyrene nanoparticles, and implement this analysis on a sample of carboxysomes, proteinaceous bacterial nanocompartments.^47,48^ Access to contrast and hydrodynamic diameter enable us to estimate single-carboxysome core-shell mass partitioning, forming the foundation for future high-precision single-particle characterization.

## 2. Results

### 2a. Trapping and interferometric signal

First, we briefly summarize the principle and implementation of the ISABEL trap, with further information provided below in Methods and in previous work.^46^ The basis of the trap is to apply active proportional feedback to a particle’s position to nearly cancel out the particle’s Brownian motion. An 808 nm CW laser is guided to the center of the ABEL microfluidic cell (Fig. 1a), steered in a Knight’s Tour laser scan pattern (Fig. 1b) by a pair of acousto-optic deflectors and detected on a dual-element balanced photoreceiver to subtract technical noise from laser power fluctuations. A field-programmable gate array (FPGA) controls the timing of the scan pattern sequence and supplies DC analog voltage feedback to bring the particle to the center of the trap (Fig. 1c). This feedback loop is updated at the end of each full tour—every 120 μs for these experiments.

**Figure 1.**
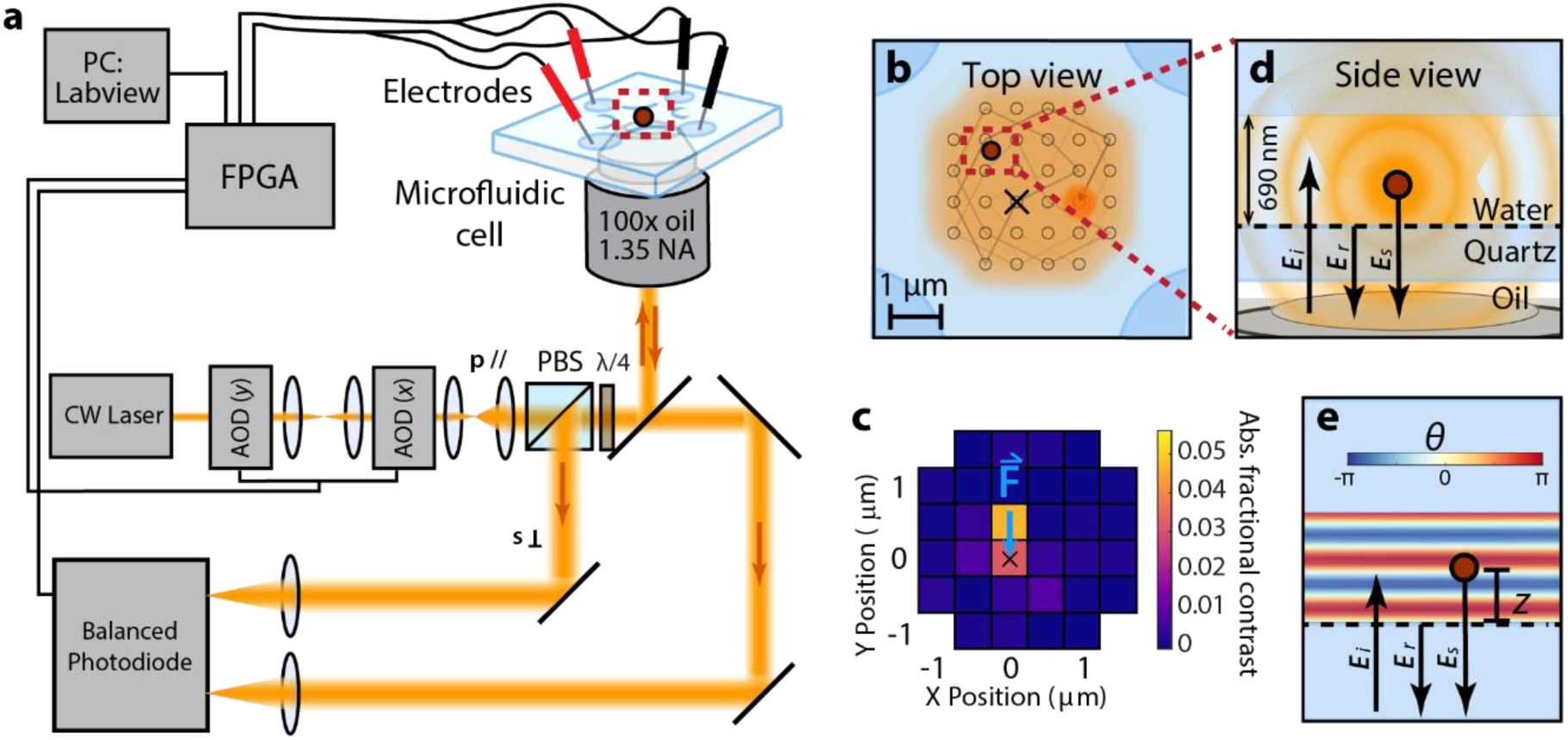
Experimental configuration of the ISABEL trap for scattering fluctuation measurements. (a) Optical scheme: a CW laser at 808 nm is scanned by a pair of acousto-optic deflectors (AODs) to form a confocal laser scan pattern in the trapping region at the center of a microfluidic cell. The interferometric scattering signal is collected on the signal arm of a balanced photodiode, isolated by a polarizing beam splitter (PBS) and a quarter-wave plate (λ/4). The reference beam for balanced detection is collected from leakage through a dichroic mirror used to overlap the scattering illumination field with a fluorescence excitation line (not shown). All analog and digital inputs and outputs are interfaced through a field-programmable gate array (FPGA) and controlled via LabVIEW. (b) Top view of the Knight’s Tour laser scan pattern in the trapping region. (c) Schematic example of the lateral feedback. The scan position exhibiting the largest absolute contrast is taken as an estimate of the particle’s position. DC electrokinetic feedback is applied so that the particle drifts to the trap setpoint in the next frame. (d) Side view of the trapping region, indicating the incident, reflected, and scattered electric fields as *E*_i_, *E*_r_, and *E*_s_, respectively. (e) Schematic of the oscillations of interferometric phase *θ* as a function of depth *z*.

Fig 1d illustrates a cross-section of the trapping region, showing the relevant optical fields for detecting interferometric scattering signal. At the sample, the incident near-IR field *E*_i_ reaches the microfluidic cell, resulting in a back-reflected field *E*_r_ and a nanoparticle scattering field *E*_s_. These fields interfere, and the total intensity is collected by the objective and sent to the signal arm of the photoreceiver. This intensity can be understood by the standard expression for interferometric scattering from a weak scatterer:^24^

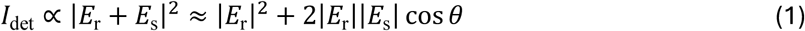

where *θ* is the relative phase between *E*_r_ and *E*_s_ and the pure scattering intensity |*E*_s_|^2^ is negligibly small. As illustrated in Fig. 1e, *θ* is in large part due to the optical path difference between the interfering *E*_r_ and *E*_s_ fields and a π/2 phase delay associated with nonresonant scattering.^31^ Thus to a first approximation, *θ*(*t*) ≈ 2*kz*(*t*) + π/2, where *k=2πn/λ* is the optical wavenumber in the liquid medium of refractive index *n,* and *z* is the axial height of the particle center from the quartz-buffer interface. This will be verified numerically in Section 2b below. We expect, then, that if the particle fluctuates in *z* due to diffusion, the phase difference *θ* will fluctuate as well. Changes in the phase *θ* are evident in the fractional scattering contrast *c*:

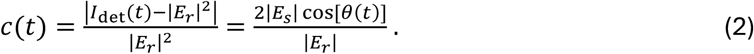

The background term |*E*_r_|^2^ is measured when feedback is turned off and thus when a particle is not trapped.

Figure 2 shows a contrast time series from trapping 100 nm polystyrene beads (data from other beads shown in Fig. S1). When the feedback is ON (white regions), the FPGA identifies the scan position with the highest absolute contrast in each full scan and applies electrokinetic feedback. Thus when a scattering particle enters the field of view, its contrast is detected and the feedback brings it to the center of the trap, as evident by the sudden jump in contrast at *t* ∼ 22 s. As described above, we see scatter contrast fluctuating between positive and negative values until the feedback is turned OFF (gray regions e.g. at *t* = 26 s) and the particle is allowed to leave the trap. To identify a trapping event and its duration for a single particle, the time trace of the absolute contrast is analyzed by a level-finding algorithm^49^ that also produces the mean absolute contrast of the particle. For these experiments, we chose the feedback ON:OFF temporal ratio of 4s:1s, which balances extensive single-particle data collection with trapping several hundred particles per hour serially. Figure 2b shows a 50-ms window of the contrast trace, highlighting the fluctuations between positive and negative values. Importantly, the contrast can linger around a particular value on the ∼ms timescale, indicating a correlation between nearby contrast values. This is distinct from white noise when the trap is empty, where sequential contrast values would be entirely random.

**Figure 2.**
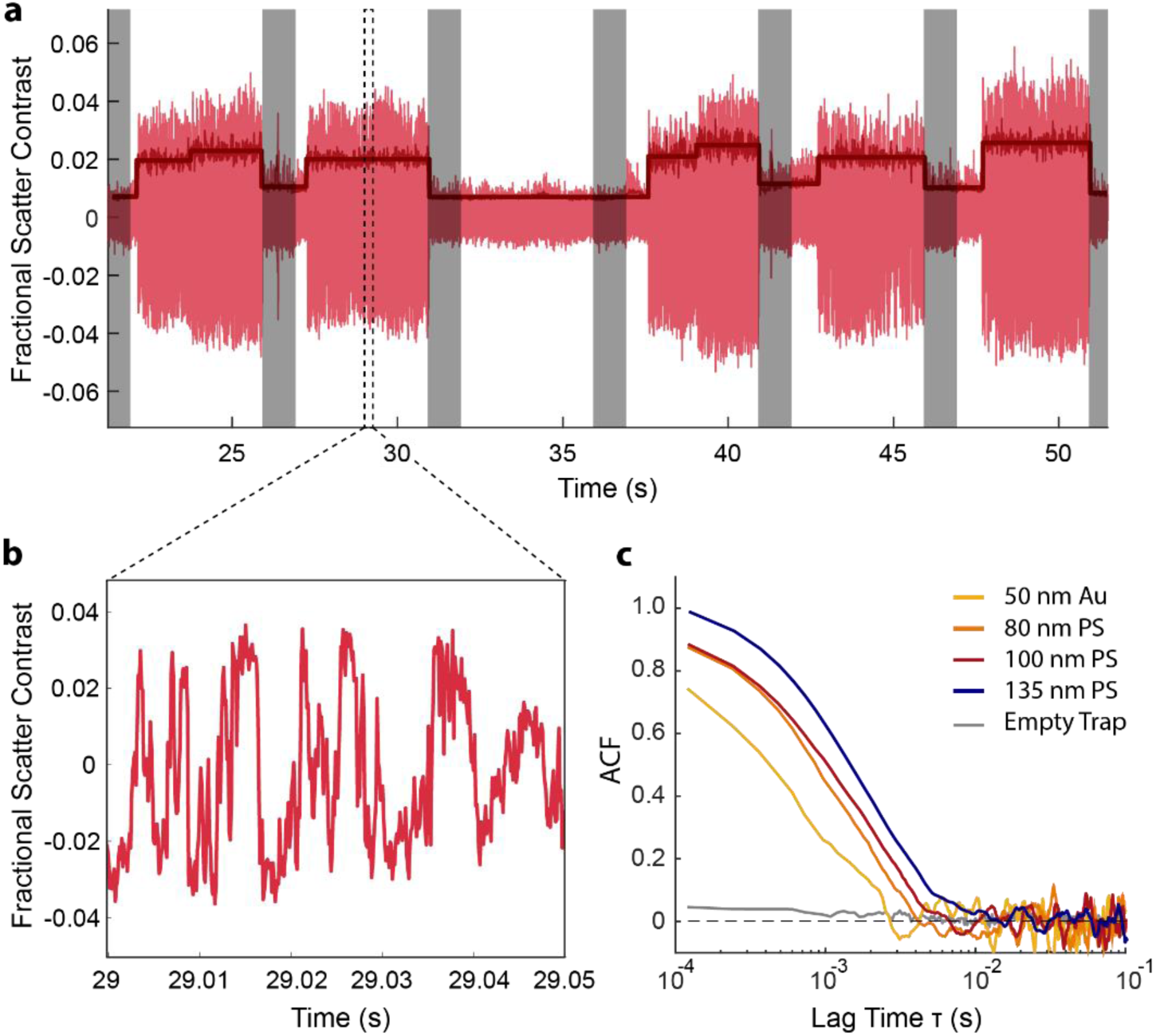
Scattering fluctuations from trapped particles. (a) An example time trace from the trap position setpoint for a sample of nominally 100nm polystyrene nanoparticles. The trace includes real-time contrast (light red), 10 ms binned absolute contrast (medium red), and absolute contrast average levels (dark red). Feedback is alternated ON:OFF for 4s:1s, where white regions are when the feedback is ON, and gray regions denote feedback OFF. (b) An enlargement of a 50-ms section of the signed scatter contrast fluctuations. (c) Time autocorrelation functions of scattering contrast from single particles representative of different samples. As nanoparticle diameter increases, the scattering contrast ACF timescale also increases.

To quantify the contrast fluctuation dynamics over each trapping event, we calculated the contrast time autocorrelation function (ACF)

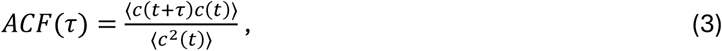

where *τ* is the time lag between two contrast measurements in a trapping event and the angled brackets refer to a temporal average. As Fig. 2c demonstrates for a typical trapped particle, contrast correlations are near 1 at short time lags (high correlation) and gradually fall to 0 (uncorrelated) with a decorrelation timescale of *τ*_1/2_ ∼ 1 ms. Importantly, Fig. 2c compares ACFs of different bead samples, spanning 50-130 nm. As particle size increases, the ACF timescale lengthens, qualitatively consistent with the inverse relationship of a particle’s diffusion coefficient and diameter according to the Stokes-Einstein equation:

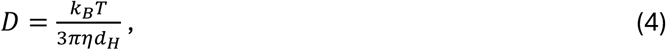

where *k*_B_ is the Boltzmann constant, *T* is the solution temperature, *η* is the solution viscosity, and *d*_H_ is the hydrodynamic diameter.

In the simple case of a Rayleigh scatterer illuminated by a plane wave and diffusing near a flat reflective surface, the contrast ACF can be solved analytically (see SI note S1) and takes the form of an exponential decay:

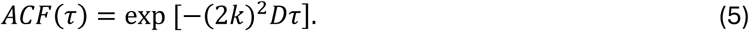

This expression demonstrates that the ACF decays more rapidly with increasing *D*, as expected. As well, the *k*-dependence of the ACF reflects that higher-frequency (i.e. shorter wavelength) light will lead to more frequent contrast change in the *z* dimension. Finally, the factor of 2 results from the doubled path length between the interfering fields due to the reflection geometry.^34,50^ Eq. 5 also arises as the special case of back-scattering in heterodyned DLS^14^ and differential dynamic microscopy.^38^

We found that our experimental ACF curves do not match Eq. 5 quantitatively (Fig S2), but rather appear stretched and shifted to longer timescales. This discrepancy arises from three key differences between our experimental configuration and the assumptions in Eq. 5: (a) Gaussian beam versus plane-wave illumination, (b) hydrodynamic confinement between two silica walls, and (c) small but non-negligible optical forces from the illumination field imparted onto the particle. To quantify each particle’s hydrodynamic diameter, we took these factors into account and assembled a simulation of nanoparticles held in the ISABEL trap.

### 2b Brownian Dynamics Simulation of the ISABEL Trap and Contrast Autocorrelation

To quantitively estimate the hydrodynamic diameter *d*_H_ and axial diffusion coefficient *D*_z_ from each trapped nanoparticle, we developed a numerical model (See Methods Section 4c below and SI Note S2). First, we simulated the three-dimensional trajectory of a particle with a predefined hydrodynamic diameter *d*_H_. This particle was allowed to diffuse without boundaries in the lateral directions *x* and *y*, but was confined in *z* within a 690 nm gap between the walls (determined experimentally by white light interferometry, SI Note S3 and SI Figs S3-S4). Simultaneously, we propagated the Knight’s Tour beam scan pattern over a 3μm × 3 μm region, completing the 32-point tour in 120 μs. The fundamental time increment in the simulation is Δ*t* = 3.75 μs, the residence time of each spot in the scan pattern. The particle undertook a new displacement step at each new Δ*t*, subject to Brownian motion, optical forces, and electrokinetic drift.

At every beam scan position, we calculated a contrast value from the particle depending on its height *z* and distance *r* from the incident beam center (SI Note S2). To efficiently implement this calculation, we first generated a contrast lookup table (Fig. 3a) based on dipole scattering near an interface.^51^ From the model, it is apparent that the contrast oscillates essentially sinusoidally as a function of *z* with period *λ/*2*n*, and decays in intensity when the particle is several hundred nanometers away from the Gaussian beam center. Given the potential contribution of lateral motion into the ACF, we assessed this possibility and found that fluctuations in *z* dominate (Fig. S5). When comparing ACFs determined from simplified contrast maps (only axial oscillations or only lateral intensity without interference), the ACF with only axial oscillations closely matches the ACF determined with the full contrast model.

**Figure 3.**
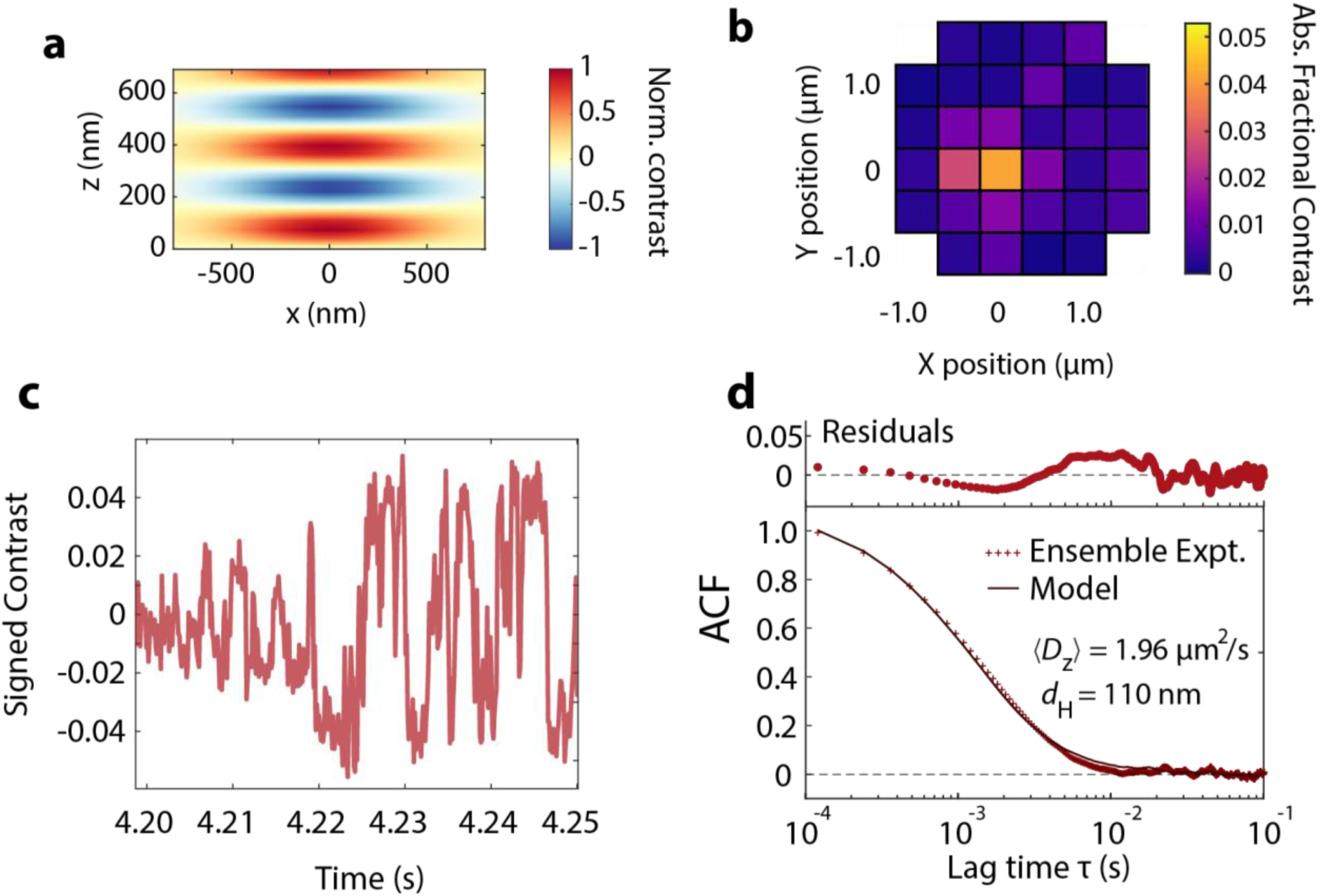
Numerical model of the ISABEL trap. (a) Contrast oscillation map for Gaussian beam illumination. The normalized interferometric contrast oscillates in *z*, as illustrated in Fig. 1e, and the amplitude of the contrast falls off with the Gaussian beam radius. (b) Example simulated frame of a trapped 100 nm polystyrene particle. (c) Example contrast time trace segment, demonstrating contrast fluctuations as seen in experiment. (d) Comparison between the ISABEL simulation and an ensemble-averaged ACF measured from 10 trapped beads. We note that the model curve was not fit to the experimental data, but parameterized according to experimental conditions (Tables S1 and S2).

At the end of each scan pattern and with contrasts determined at each point, the scan position with the highest absolute contrast was taken as the estimate of the particle position for the purposes of applying feedback. An example frame of a simulated 100 nm polystyrene bead is shown in Fig 3b, where we added Gaussian white noise to each scan position consistent with experiment. The particle’s response to the feedback was parameterized with an effective electrokinetic mobility to ensure reliable trapping in the model. Over the next iteration of the Knight’s Tour, DC electrokinetic drift imparts a constant particle velocity until a new feedback voltage is applied.

We also modeled *z*-dependent anisotropic diffusion to account for the effects of hydrodynamic drag near the walls. Because long-range collective forces acting on the particle are suppressed near a solid boundary, nanoparticles are known to experience reduced diffusion as they approach an interface.^52,53^ The effect of this is well approximated by a closed-form expression (SI Note S2), and breaks the symmetry of the diffusion coefficient such that *D*_z_ perpendicular to the wall is attenuated more than *D*_xy_ (Fig. S6). Following observations from previous experiments,^53^ we assume the effects from two walls independently act on the particle. The results from this treatment provide us with *z*-dependent diffusion coefficients *D*_z_(*z*) and *D*_xy_(*z*), updated every Δ*t*.

Finally, our model accounts for a weak scattering force biasing the nanoparticle to the top of trapping region. In our experiments, we sent ∼20 mW to the sample to generate optimal photocurrent on our photodetector from the few-percent quartz-water reflection, leading to a time-averaged intensity of 16 kW/cm^2^ over the scan pattern. To account for the impact of this irradiation, we incorporated optical scattering and gradient forces into our simulation (SI note S2), since we saw that confinement effects were still insufficient to capture the experimental ACF (Fig. S2). Even though lateral gradient forces turned out to be negligible, a weak scattering force biased particles towards higher regions near the top wall and into a region of attenuated diffusion. Additionally, we found that this bias explains observed asymmetric histograms of the contrast values from a trapping event (Fig. S7), supporting the idea that the particle sampled *z* unevenly during trapping. Without this effect, the contrast histogram would be symmetric about zero, inconsistent with our observations.

The additional considerations of Gaussian illumination, confined diffusion, and optical forces enable a simulation of scattering contrast fluctuations (Fig. 3c) consistent with experiment. More quantitatively, we observe remarkable agreement with a model ACF to experimental ACFs (Fig. 3d). As illustrated here, we refer to the extracted diffusion coefficient of the confined particle as 〈*D*_z_〉 to reflect the temporal average of *D*_z_ values experienced as the particle explores the microfluidic depth. The satisfactory agreement between model and experiment is maintained across four standard bead samples (SI Fig. S8), where new simulations were generated for different classes of particles and their scattering cross-sections (see Note S2 and Table S2). With this model in hand, we could evaluate the ACFs from individual beads.

### 2c Sizing individual beads with model ACFs

From our ACF model, we developed a library of simulated ACF curves covering a distribution of hydrodynamic diameters and materials, to be used as reference for experimental ACFs (see Table S2 for particle-specific parameters). For each experimental ACF, we implemented a weighted least-squares fit to each of the reference ACF curves to determine the best model for the data. An example result from this optimization is shown in Fig. 4a, where we retrieve a 〈*D*_z_〉 = 1.87 ± 0.22 μm^2^/s (mean ± standard error of the mean, SEM) and *d*_H_ = 113 ± 4 nm from a bead sampled from the population of nominally 100 nm polystyrene beads. To assess the uncertainty on the measurement and analysis procedure, we divided each experimental trapping event into three sub-intervals and calculated the SEM from the subdivided measurements. Uncertainty decreases with longer trapping time (SI Fig. S9) so that with 4s of trapping, median 〈*D*_z_〉 relative uncertainty is 8% and median *d*_H_ uncertainty is 4%.

**Figure 4.**
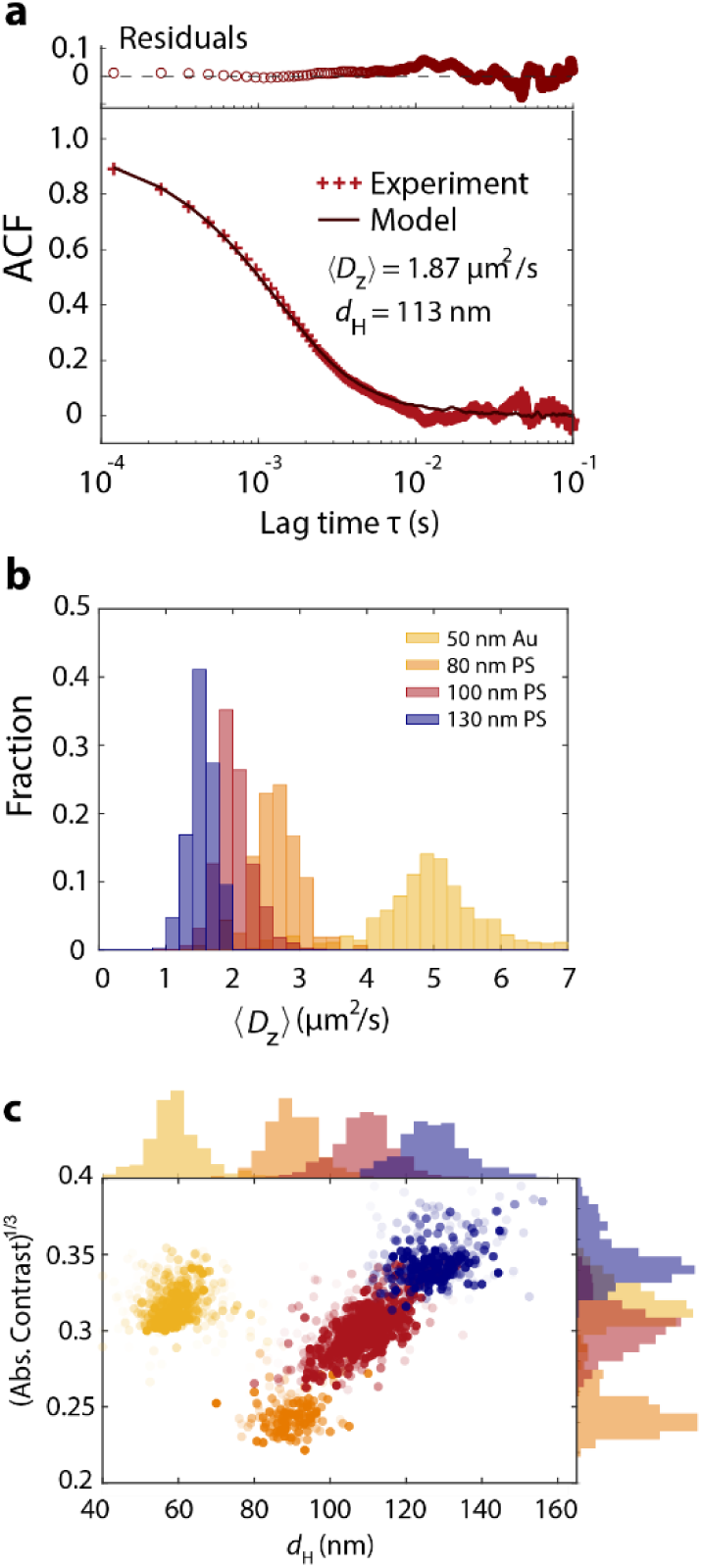
Statistics of estimated diffusion coefficients and hydrodynamic diameters. (a) Example experimental ACF and model fit. (b) Histograms of extracted diffusion coefficients 〈*D_z_*〉 for the four standard bead samples. (c) Two-dimensional scatter plot and marginal histograms for contrast and extracted hydrodynamic diameter. The color scheme is the same as in (a). Opacity of each point indicates trapping length, from 0.2-4.0s (lightest to darkest).

Our estimated 〈*D*_z_〉 and *d*_H_ distributions over standard particle samples are displayed in Fig. 4b and 4c, respectively. For the polystyrene beads where 〈*D*_z_〉 < 3 μm^2^/s, we observe narrow distributions consistent with expectations based on the low polydispersity from these samples. However, we find that the precision on 〈*D*_z_〉 is lowest for 50 nm Au beads. Because ACF from 50 nm gold decays most rapidly, the early-time plateau of the ACF is not captured within the time resolution of our measurement (see Fig. S8c). This sample is the most monodisperse (σ/μ = 2% by TEM), implying an expected 〈*D*_z_〉 standard deviation of ∼0.1 μm^2^/s according to Eq 4. However, the extracted 〈*D*_z_〉 distribution displays significantly more spread with *σ* = 0.88 μm^2^/s. Analyzing the ACFs from 100 identically simulated beads of *d*_H_ = 59 nm yields *d*_H_ = 58.6 ± 3.5 nm (μ ± σ) and 〈*D*_z_〉 = 4.95 ± 0.39 μm^2^/s (Fig. S10). The extra variability retrieved from experiment suggests additional sources of uncertainty in experiment, such as residual optical noise and limited time resolution.

Fig 4c shows the 2D scatter plot comparing extracted *d*_H_ values and absolute contrast for each particle. We present the cube root of contrast, since contrast is expected to scale linearly with the particle volume for uniform objects.^23^ For the population of 100 nm polystyrene beads, we observe a linear trend between contrast^1/3^ and *d*_H_ within the sample, and we also see 80 nm and 130 nm polystyrene samples on the same trendline. Table 1 shows that the median *d*_H_ values for each standard sample are within agreement of expectations from DLS and TEM measurements, with some deviation in the 80 nm polystyrene sample. We note that we expect *d*_H_ to be a few nm larger than TEM diameters, but the differences from this expectation are generally smaller than the uncertainty of our measurement. Encouragingly, the 50 nm AuNP sample exhibits a slight positive correlation between contrast^1/3^ and *d*_H_, despite the sample’s monodispersity, highlighting the high sensitivity of interferometric contrast to particle volume. With the particle sizing validated by standard samples, we tested our sizing procedure on a multi-component biological nanoparticle: the carboxysome.

**Table 1:** Extracted and reference sizing statistics for trapped nanoparticles.

| **Nanoparticle** | ***N*** | **$D_z \mu \pm \sigma$**<br>**($\mu\text{m}^2/\text{s}$)** | **Median**<br>**SEM on $D_z$**<br>**($\mu\text{m}^2/\text{s}$)** | **$d_H \mu \pm \sigma$**<br>**(nm)** | **$d \mu \pm \sigma$ (nm)**<br>**Reference**<br>**(method)** | **Median**<br>**SEM on $d_H$**<br>**(nm)** |
| --- | --- | --- | --- | --- | --- | --- |
| 50 nm Au | 491 | $4.96 \pm 0.88$ | 0.37 | $59.4 \pm 7.3$ | $59 \pm 1.9$<br>(DLS) | 2.7 |
| 80 nm PS | 161 | $2.66 \pm 0.36$ | 0.18 | $90.0 \pm 6.4$ | $81 \pm 9.5$<br>(PCS) | 3.3 |
| 100nm PS | 671 | $2.02 \pm 0.28$ | 0.15 | $108.8 \pm 8.1$ | $108 \pm 7$<br>(TEM) | 4.4 |
| 130 nm PS | 314 | $1.54 \pm 0.20$ | 0.12 | $127.5 \pm 9.0$ | $130 \pm 2.4$<br>(TEM) | 5.5 |
| carboxysomes | 549 | $2.04 \pm 0.64$ | 0.40 | $117.3 \pm 24.8$ | $112 \pm 17$<br>(TEM) | 13.3 |

### 2d Sizing and core mass of carboxysomes

Carboxysomes are a fascinating example of bacterial microcompartments, responsible for 10-25% of global carbon fixation^48^ and an important “proto-organelle” in various autotrophic bacteria.^47,54^ The compartment consists of a proteinaceous shell, internally loaded enzymes carbonic anhydrase and Rubisco, and the disordered scaffolding protein CsoS2 (Fig. 5a-b). Recent mutagenesis studies revealed the role of CsoS2 in binding cargo and affecting carboxysome diameter,^55–57^ though it is also expected that net enzymatic kinetics would be influenced by cargo packing density. This variable cargo packing density also means that scattering contrast is not a reliable proxy for particle shell size. To test the efficacy and generality of our sizing procedure, we reanalyzed past carboxysome trapping data where we used interferometric scattering contrast to estimate the mass of each carboxysome.^45^

**Figure 5.**
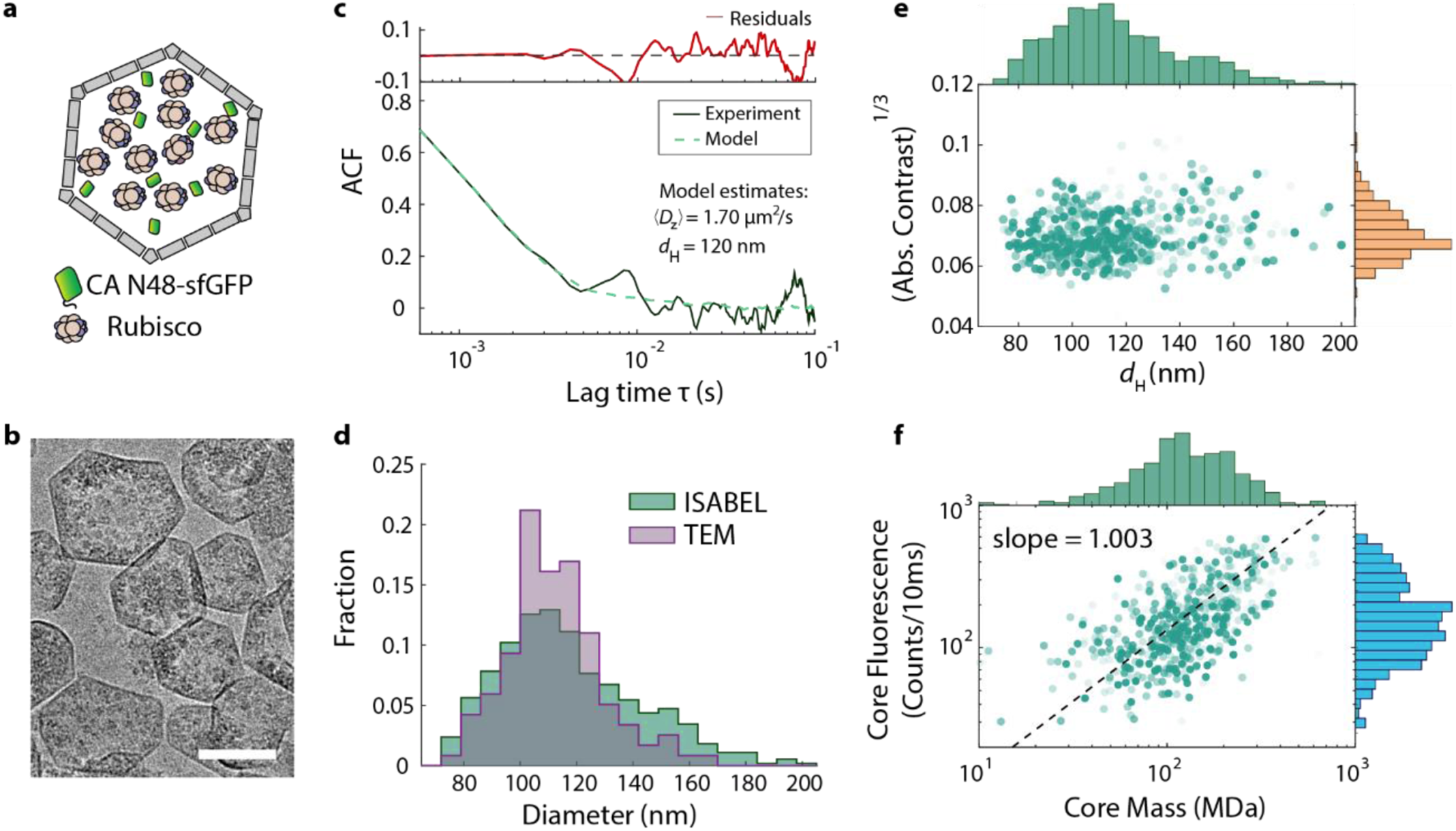
Size and mass partitioning from individual carboxysomes in the ISABEL trap. (a) Schematic of the carboxysome. Carboxysomes contain the enzyme Rubisco surrounded by a proteinaceous shell. We examined carboxysomes that were additionally loaded with sfGFP with a sequence that targets binding to Rubisco. (b) CryoTEM image of carboxysomes. Scale bar 100 nm. (c) A representative experimental and model ACF from a trapped carboxysome. (d) Histograms of hydrodynamic diameters from the ISABEL trap, and effective diameters measured from TEM. (e) Two-dimensional scatter plot and marginal histograms for contrast and extracted hydrodynamic diameter. Opacity of each point indicates trapping length, from 0.2-4.0s (lightest to darkest). (f) Core mass results from core-shell Mie scattering model, incorporating new diameter estimates. Core mass scales linearly with core-loaded fluorescence, in agreement with expectation. Panels (a) and (b) reproduced with permission from The American Chemical Society, copyright 2022.

To analyze this data, we generated a new set of reference ACF curves to match past experimental conditions (see Table S2). An example of a typical carboxysome ACF is shown in Fig. 5c. From our past TEM measurements (Table 1 and Fig. 5d), we expect the median *d*_H_ ∼ 110 nm leading to a lower 〈*D*_z_〉, which could still be estimated with reasonable precision despite the lower time resolution. Even though the ACF curve at high correlations is not recovered (Fig. 5c), we find satisfactory agreement with TEM diameters (Fig. 5d). Distribution peaks are comparable between TEM and ISABEL, though there is some excess spread and a high-diameter shoulder around 150 nm in ISABEL data, which we attribute to a known subpopulation of carboxysome dimers. The resulting contrast^1/3^-*d*_H_ scatter plot is shown in Fig. 5e, revealing a weak positive correlation, less well defined than in the standard samples. The subpopulation with higher diameters is less geometrically regular and more likely to release cargo, which may explain the unexpectedly low contrast.

Finally, we used the newly available diameter information to revisit our previous model for cargo loading, where before we relied on the assumption that fluorescence brightness from internalized sfGFP linearly scaled with core mass. Now, with direct diameter measurement for each carboxysome, we could re-employ our Mie-scattering core-shell model^45^ to determine the core refractive index, and therefore mass of core protein. Fig. 5f shows the two-dimensional scatter plot comparing retrieved core mass with the measured core fluorescence brightness, empirically demonstrating direct correlation. By fitting the scatter plot to a power law, we get a remarkably accurate slope of 1.003, in strong agreement with our previous assumption. This measurement capability, in combination with diffusion sizing information, allows us to accurately analyze core-shell mass partitioning, which frees up the fluorescence channel for distinct spectroscopic reporting.

## 3. Discussion and Conclusion

This study demonstrates the effectiveness of sizing individual nanoparticles by analyzing interferometric scattering fluctuations. We find that detailed knowledge of experimental parameters is necessary to obtain accurate *d*_H_ values, though this is not unusual for single-particle tracking measurements.^22,58^ Due to the high *z* sensitivity of interferometric scattering signal and the relatively coarse 500 nm spacing of the Knight’s Tour pattern, this sizing method displays distinct benefits compared to tracking lateral displacements of a nanoparticle in our trap. A useful feature of the ACF analysis is that residual white noise does not impact the accuracy of the ACF, even if the noise impacts the precision of the measurement. These trapping measurements enabled substantial data collection (up to 32,000 points per particle) due to trapping for up to 4s. Precision on a particle’s *d*_H_ estimate could be further improved below our achieved 3-5 nm precision with longer trapping, improved time resolution, or reduction in microfluidic height to accentuate differences in diffusion attenuation. We note that while our methods yielded satisfactory accuracy, other studies of nanoparticles in nanofluidic channels have observed anomalously low diffusion (even accounting for hydrodynamic effects), though those studies were generally in narrower channels (<500 nm) and with much lower ionic strengths (< 1 mM) than we employed.^59–63^

Our study, while not the first to size particles by interferometric scattering fluctuations, provides important lessons applicable to other experiments. First, Eq. 5 highlights the method’s sensitivity to not just *D*, but also the wavenumber of the incident light. Indeed, even with 808 nm excitation, we found imprecision in sizing 50 nm Au particles, despite strong contrast signal. For a change of illumination to the commonly used 445 nm, the ACF would decay ∼3× faster. Thus for a particular nanoparticle sample, a trade-off emerges between enhanced contrast and faster ACF timescales (and thus required sampling rates) with shorter wavelength illumination. Optical effects such as illumination geometry and intensity will also play a role in the resulting contrast fluctuations. Careful consideration of each of these parameters equips the experimenter with design principles for label-free measurements of particle dynamics.

In summary, we demonstrated that individual nanoparticles in an anti-Brownian electrokinetic trap can be sized using a statistical analysis of scattering fluctuations. As with other interferometric scattering measurements, sizing information is readily obtainable label-free from both contrast and hydrodynamic fluctuation, with the combination allowing detailed study of nanoparticles of variable density. To our benefit, trapping extends observation of a single particle in free solution, reliably providing thousands of contrast data points per particle. With interpretive aid from a Brownian Dynamics model of the ISABEL trap, we could quantify individual particles’ hydrodynamic diameters from the contrast time ACF. Standard gold and polystyrene nanoparticles benchmarked the accuracy and precision of the method, allowing us to also estimate the method’s performance on bacterial carboxysome nanocompartments. The combination of contrast and diameter allowed us to take these measurements a step further to validate a direct relationship between core mass and core fluorescence. By leveraging interferometric scattering for these single-particle nanoscale characteristics, we can reserve fluorescence channels for spectroscopic measurements, as we have demonstrated before.^46^ Looking forward, the combination of label-free nanoparticle characterization with chemical dynamics measurements afforded by fluorescence spectroscopy will reveal never-before-seen richness and detail in nanoparticle measurement.

## 4. Methods

### 4a Materials and microfluidic preparation

Nanoparticle samples were prepared according to previously described procedures,^45^ summarized here. Carboxyl-PEG 50 nm UltraUniform Gold Nanoparticles were purchased from NanoComposix (Cat. No. AUXU50, Lot No. SDC0158). Carboxylated 100 nm polystyrene Fluospheres (505/515 fluorescence excitation/emission) were purchased from ThermoFisher Scientific (Cat. No. F8803, Lot no. 1588588). Carboxylated 130 nm polystyrene nanoparticles were acquired from Bangs Laboratories (Cat. No. PC02005, Lot no. 15033). Carboxylated, NIST-traceable 80 nm polystyrene nanoparticles were obtained from ThermoFisher Scientific (Cat. No. 3080A, lot no. 242108) with certified mean diameter of 81 ± 3 nm, (95% confidence interval). Carboxysomes were purified from *E. coli* as previously described.^45^

Before each experiment, we passivated the ABEL microfluidic cell with a polyelectrolyte multilayer consisting of four alternating layers of cationic poly(ethyleneimine) and anionic poly(acrylic acid) to provide an anionic surface that electrostatically repelled particles from sticking to the microfluidic walls. The same microfluidic cell was used for all experiments of standard beads, piranha cleaned and freshly passivated for each experiment, to ensure the same microfluidic cell height across experiments. All nanoparticle samples were taken from stock solutions and diluted to single-particle concentrations (∼10 pM) in a buffer at pH 7.5 consisting of 10 mM HEPES and 15 mM NaCl, and sonicated for 10 minutes before introduction into the passivated microfluidic cell.

### 4b Optical setup

We summarize the implementation of the ISABEL trap, with further information provided in previous work.^46^ Illumination for scattering signal was provided by a multimode 808 nm laser diode (Qphotonics QFLD-808-250S), driven by a low-noise laser diode driver (Koheron CTL-200-1-B-600), which includes thermoelectric cooling, a bias tee for RF diode modulation, and USB port for serial communication. To reduce the beam coherence, a bias tee in the diode mount was supplied with a sinusoidal AC wave from a function generator (Agilent 33220A) at 2.5MHz with V_pk-pk_ = 5.00 V. With 50Ω impedance, this corresponds to a current modulation of 100 mA peak-peak on the 300 mA DC current from the laser diode driver. The beam was initially horizontally (*p*) polarized, then rotated to circular polarization with a zero-order quarter-wave plate before entering a 100x, 1.35 NA oil immersion objective (Olympus UplanApo). When the back-reflected beam and scattered beams propagated in reverse out of the objective lens, they rotated to vertical polarization via the quarter-wave plate and isolated from the incoming beam via a polarizing beam splitter. The signal was collected on the signal arm of a balanced photodiode (New Focus 1807). The balanced photodiode was introduced to correct for residual laser power fluctuations, using a reference beam that is separated just before the objective via the dichroic for overlapping with a fluorescence excitation line.

To generate a two-dimensional (x-y) map in the sample plane with point detection, the beam is scanned in a deterministic Knight’s Tour pattern by a pair of acousto-optic deflectors (AA Optoelectronics MT110-B50A1,5-IR, driven by AA Opto Electronic DDSPA2X-D4125b-34).^64^ The result is a confocal beam scanning pattern in the sample plane, comprised of 32 positions spaced 500nm apart and completed in 120 μs. The photodiode signal is sent to a National Instruments field-programmable gate array (FPGA, R-7852) to synchronize signal detection with the deterministic scan pattern. A feedback look-up table determines the DC analog voltage feedback, proportional to the distance between the beam scan position with largest absolute contrast and the feedback setpoint.

Determining contrast required a real-time estimation of background reflection. The background term |*E*_r_|^2^ was taken as mean signal over 128 frames when feedback was turned off. Though contrast was determined in real-time, the raw *I*_det_ was saved for post-processing to recover the signed contrast.

### 4c Trapping simulations and single-particle diameter determination

We constructed a Brownian Dynamics model of the ISABEL trap for comparison to experimental contrast autocorrelation functions. In summary, the particle with diameter *d*_H_ was allowed to diffuse at every time step Δ*t* = 3.75 μs (Table S2), physically confined in the axial direction but unconfined in the lateral dimensions. Upon reaching the upper or lower boundary in the axial direction, the particle was reflected according to hard-sphere collision.

At each time step, the beam was stepped to a new position in the 32-point Knight’s Tour pattern.^64^ Based on the location of the particle with respect to the beam center, interferometric contrast and optical forces were calculated at each scan position, as described in SI Note S2. At the completion of a Knight’s Tour, the position of the absolute maximum contrast *x*_max_ was used as the estimate of the particle’s position. Based on the feedback gain *g* given in Table S1, voltage *V*(*t*) = - *gx*_max_(*t*) and particle electrokinetic mobility *μ*, an electrokinetic drift *v*(*t*) = *μV*(*t*) was applied to the particle’s trajectory over the course of the next Knight’s tour.

Overall, the particle trajectory was propagated according to the following equations:

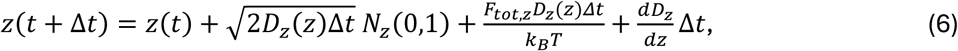

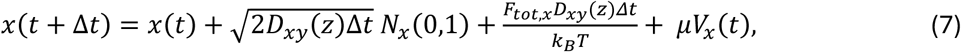

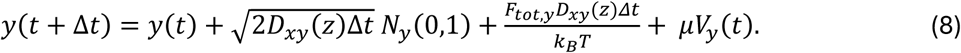

The second terms on the right-hand side represent the Brownian displacement characterized by diffusion coefficient *D*(*z*) and normal random variable *N*(0,1) with μ = 0 and σ = 1. As described further in the SI Note S2, we calculate *D*_xy_(*z*) and *D*_z_(*z*) at each time step due to variations in hydrodynamic forces as a function of distance from the microfluidic walls. The third righthand terms represent the displacements due to optical forces applied to the particle, also explained further SI Note S2. The last righthand term in Eq. 6 reflects particle translation due to the gradient in *z* of the diffusion coefficient.^65^ To approximate this, we determined the diffusion coefficients ±1 nm flanking the particle’s position and calculated the average of the finite differences Δ*D*_z_(*z*)/Δ*z*. Finally, the fourth terms in Eqs. 7 and 8 are the applied electrokinetic drift.

We used the above model to generate contrast fluctuation trajectories for assembly of a library of contrast ACFs. For each class of particle, we simulated the trapping a single particle for 80s, repeating the simulation for a range of *d*_H_ values spanning 50 nm in increments of 1 nm, centered on the nominal *d*_H_ value. For Au 50 nm particles, we sampled the *d*_H_ range in 0.5 nm steps. To estimate the *d*_H_ from an experimental ACF, we calculated the root-mean-square error (RMSE) between the experimental and model curves, scanning the space of *d*_H_ and ACF amplitude, which was generally less than 1 due to measurement white noise. The model ACF with the lowest RMSE was selected for an estimate of the particle’s *d*_H_ and *D*_z_. To capture ∼90% of the ACF drop without excess influence from noise at long times, we additionally only implemented the fit for lag time *τ* < 0.003 s for the polystyrene samples and *τ* < 0.0015 s for the 50 nm Au beads.

To assess uncertainty from each ACF diameter assignment, we subdivided the contrast trajectory into thirds, implemented the fit separately, and calculated the mean and standard error of the mean (SEM) = *σ*/√3. SEMs on estimated *d*_H_ and *D*_z_ are shown in Fig. S9, which demonstrate improved precision with increased trapping time.

## Acknowledgments

This work was supported by Stanford University and The Ohio State University. We thank Prof. W.E. Moerner for valuable discussions and for use of apparatus. Finally, we acknowledge Dr. Luke Oltrogge and Prof. David Savage for insightful comments and for providing carboxysome sample.

## Data Availability

The data that support the findings of this study can be obtained from the corresponding author upon request.

## Author Contributions

Conceptualization, AAL and WBC. Methodology, WBC; Investigation, WBC; Software, AAL and WBC; Visualization, WBC; Writing – Original Draft, WBC; Writing – Review and Editing, AAL and WBC; Resources, WBC; Project Administration, WBC.

## References

1 P. Zijlstra, P. M. R. Paulo and M. Orrit, Nat. Nanotechnol., 2012, 7, 379–382.

2 M. J. Mitchell, M. M. Billingsley, R. M. Haley, M. E. Wechsler, N. A. Peppas and R. Langer, Nat. Rev. Drug. Disc., 2021, 20, 101–124.

3 H. Kirst, B. H. Ferlez, S. N. Lindner, C. A. R. Cotton, A. Bar-Even and C. A. Kerfeld, Proc. Natl. Acad. Sci. U. S. A., 2022, 119, e2116871119.

4 D. Gao, H. Zhou, J. Wang, S. Miao, F. Yang, G. Wang, J. Wang and X. Bao, J. Am. Chem. Soc., 2015, 137, 4288–4291.

5 B. Y. Shekunov, P. Chattopadhyay, H. H. Y. Tong and A. H. L. Chow, Pharm. Res., 2007, 24, 203–227.

6 W. Jiang, B. Y. S. Kim, J. T. Rutka and W. C. W. Chan, Nat. Nanotechnol., 2008, 3, 145–150.

7 A. A. Koelmans, E. Besseling and W. J. Shim, in Marine anthropogenic litter, ed. Bergmann, Melanie, L. Gutow, Klages and Michael, Springer International Publishing, 2015, p. 325–340.

8 K. Tiede, A. B. A. Boxall, S. P. Tear, J. Lewis, H. David and M. Hassellöv, Food Addit. Contam. - Chem. Anal. Control Expo. Risk Assess., 2008, 25, 795–821.

9 S. M. Stavis, J. A. Fagan, M. Stopa and J. A. Liddle, ACS Appl. Nano Mater., 2018, 1, 4358–4385.

10 E. Patois, M. A. H. Capelle, C. Palais, R. Gurny and T. Arvinte, J. Drug. Deliv. Sci. Technol., 2012, 22, 427–433.

11 B. Huang, M. Bates and X. Zhuang, Ann. Rev. Biochem., 78, 993–1016.

12 J. Schnitzbauer, M. T. Strauss, T. Schlichthaerle, F. Schueder and R. Jungmann, Nat. Protoc., 2017, 12, 1198–1228.

13 P. Kondylis, C. J. Schlicksup, A. Zlotnick and S. C. Jacobson, Anal. Chem., 2019, 91, 622–636.

14 B. J. Berne and R. Pecora, Dynamic Light Scattering with Applications to Chemistry, Biology, and Physics, Dover, 2000.

15 M. A. Digman and E. Gratton, Ann. Rev. Phys. Chem., 2011, 62, 645–668.

16 L. Yu, Y. Lei, Y. Ma, M. Liu, J. Zheng, D. Dan and P. Gao, Front. Phys., 2021, 9.

17 J. Gallego-Urrea, J. Tuoriniemi and M. Hassellöv, Trends Anal. Chem., 2011, 30, 473–483.

18 A. C. Madison, A. L. Pintar, C. R. Copeland, N. Farkas and S. M. Stavis, ACS Nano, 2023, 17, 8837–8842.

19 V. Filipe, A. Hawe and W. Jiskoot, Pharm. Res., 2010, 27, 796–810.

20 B. Špačková, H. Klein Moberg, J. Fritzsche, J. Tenghamn, G. Sjösten, H. Šípová-Jungová, D. Albinsson, Q. Lubart, D. van Leeuwen, F. Westerlund, et al, Nat. Meth., 2022, 19, 751–758.

21 A. D. Kashkanova, M. Blessing, M. Reischke, J. Baur, A. S. Baur, V. Sandoghdar and J. Van Deun, J Extracell Vesicles., 2023, 12, 12348.

22 A. D. Kashkanova, M. Blessing, A. Gemeinhardt, D. Soulat and V. Sandoghdar, Nat. Meth., 2022, 19, 586–593.

23 R. W. Taylor and V. Sandoghdar, Nano Lett., 2019, 19, 4827–4835.

24 N. S. Ginsberg, C. Hsieh, P. Kukura, M. Piliarik and V. Sandoghdar, Nat. Rev. Methods Primers, 2025, 5, 23.

25 G. Young and P. Kukura, Ann. Rev. Phys. Chem., 2019, 70, 301–322.

26 S. Spindler, J. Ehrig, K. König, T. Nowak, M. Piliarik, H. E. Stein, R. W. Taylor, E. Garanger, S. Lecommandoux, I. D. Alves, et al, J. Phys. D, 2016, 49, 274002.

27 B. Wu, S. Tsai and C. Hsieh, J. Phys. Chem. C, 2025, 129, 5075–5085.

28 M. Küppers, D. Albrecht, A. D. Kashkanova, J. Lühr and V. Sandoghdar, Nat. Commun., 2023, 14, 1962.

29 G. Young, N. Hundt, D. Cole, A. Fineberg, J. Andrecka, A. Tyler, A. Olerinyova, A. Ansari, E. G. Marklund, M. P. Collier, et al, Science, 2018, 360, 423–427.

30 A. D. Kashkanova, D. Albrecht, M. Küppers, M. Blessing and V. Sandoghdar, ACS Nano, 2024, 18, 19161–19168.

31 R. Gholami Mahmoodabadi, R. W. Taylor, M. Kaller, S. Spindler, M. Mazaheri, K. Kasaian and V. Sandoghdar, Opt. Express, 2020, 28, 25969–25988.

32 B. van Heerden and T. P. J. Krüger, J. Chem. Phys., 2022, 157, 084111.

33 K. Kasaian, M. Mazaheri and V. Sandoghdar, ACS Nano, 2024, 18, 30463–30472.

34 Y. Hsiao, I. Liao, B. Wu, H. C. Chu and C. Hsieh, Commun. Biol., 2024, 7, 763.

35 L. Kisley, R. Brunetti, L. J. Tauzin, B. Shuang, X. Yi, A. W. Kirkeminde, D. A. Higgins, S. Weiss and C. F. Landes, ACS Nano, 2015, 9, 9158–9166.

36 L. Needham, C. Saavedra, J. K. Rasch, D. Sole-Barber, B. S. Schweitzer, A. J. Fairhall, C. H. Vollbrecht, S. Wan, Y. Podorova, A. J. Bergsten, et al, Nature, 2024, 629, 1062–1068.

37 S. Palkhivala, L. Kohler, C. Ritschel, C. Feldmann and D. Hunger, ACS Nano, 2025,.

38 R. Cerbino, F. Giavazzi and M. E. Helgeson, J. Polym. Sci., 2022, 60, 1079–1089.

39 K. A. Ibrahim, A. S. Naidu, H. Miljkovic, A. Radenovic and W. Yang, ACS Nano, 2024, 18, 10738–10757.

40 W. B. Carpenter, A. A. Lavania, A. H. Squires and W. E. Moerner, J. Phys. Chem. C, 2024, 128, 20275–20286.

41 A. H. Squires, A. A. Lavania, P. D. Dahlberg and W. E. Moerner, Nano Lett., 2019, 19, 4112–4117.

42 A. H. Squires, A. E. Cohen and W. E. Moerner, Anti-Brownian Traps, Springer Berlin Heidelberg, 2018.

43 Q. Wang and W. E. Moerner, Nat. Meth., 2014, 11, 555–558.

44 A. E. Cohen and W. E. Moerner, Appl. Phys. Lett., 2005, 86, 093109.

45 A. A. Lavania, W. B. Carpenter, L. M. Oltrogge, D. Perez, J. B. Turnšek, D. F. Savage and W. E. Moerner, J. Phys. Chem. B, 2022, 126, 8747–8759.

46 W. B. Carpenter, A. A. Lavania, J. S. Borden, L. M. Oltrogge, D. Perez, P. D. Dahlberg, D. F. Savage and W. E. Moerner, J. Phys. Chem. Lett., 2022, 13, 4455–4462.

47 T. O. Yeates, C. A. Kerfeld, S. Heinhorst, G. C. Cannon and J. M. Shively, Nat. Rev. Microbiol., 2008, 6, 681–691.

48 J. S. Borden and D. F. Savage, Curr. Opin. Microbiol., 2021, 61, 58–66.

49 S. Yin, N. Song and H. Yang, Biophys. J., 2018, 115, 217–229.

50 Y. Hsiao, C. Tsai, T. Chen and C. Hsieh, ACS Nano, 2022, 16, 2774–2788.

51 A. S. Backer and W. E. Moerner, J. Phys. Chem. B, 2014, 118, 8313–8329.

52 X. Bian, C. Kim and G. E. Karniadakis, Soft Matter, 2016, 12, 6331–6346.

53 B. Lin, J. Yu and S. A. Rice, Phys. Rev. E, 2000, 62, 3909–3919.

54 C. A. Kerfeld and M. R. Melnicki, Curr. Opin. Plant Biol., 2016, 31.

55 L. M. Oltrogge, A. W. Chen, T. Chaijarasphong, J. B. Turnšek and D. F. Savage, Biochemistry, 2024, 63, 219–229.

56 T. Ni, Q. Jiang, P. C. Ng, J. Shen, H. Dou, Y. Zhu, J. Radecke, G. F. Dykes, F. Huang and L. Liu, Nat. Commun., 2023, 14, 5512.

57 L. Tianpei, C. Taiyu, C. Ping, G. Xingwu, C. Vincent, Dykes Gregory F., W. Qiang and Liu Lu-Ning, mBio, 2024, 15, 1358.

58 X. Michalet and A. J. Berglund, Phys. Rev. E, 2012, 85, 061916.

59 A. Behjatian, M. Bespalova, N. Karedla and M. Krishnan, Phys. Rev. E, 2020, 102, 042607.

60 N. Mojarad and M. Krishnan, Nat. Nanotechnol., 2012, 7, 448–452.

61 H. T. Hoang, I. Segers-Nolten, N. R. Tas, J. W. van Honschoten, V. Subramaniam and M. C. Elwenspoek, Nanotechnology, 2011, 22, 275201.

62 S. L. Eichmann and M. A. Bevan, Langmuir, 2010, 26, 14409–14413.

63 Y. Kazoe, K. Mawatari and T. Kitamori, Anal. Chem., 2015, 87, 4087–4091.

64 Q. Wang and W. E. Moerner, Appl. Phys. B, 2010, 99, 23–30.

65 D. L. Ermak and J. A. McCammon, J. Chem. Phys., 1978, 69, 1352–1360.

