## Supplemental Information for "Sizing single trapped nanoparticles with interferometric scattering fluctuations"

### Note S1: Derivation of analytical contrast autocorrelation function

Consider a Rayleigh scatterer diffusing above a reflective interface according to an isotropic diffusion coefficient *D* and illuminated by a plane wave *E*cos(*kz*+ωt). For simplicity, we assume monochromatic illumination and infinite coherence length. Because of the optical delay between the reflected and scattered fields, the interferometric scattering contrast can be described as:

$$c(t) = \frac{2|E_s|\cos[2kz(t)]}{|E_r|} \propto \cos[2kz(t)] \quad \text{(S1)}$$

Therefore the time autocorrelation function of the contrast can be written down as

$$ACF(\tau) = \frac{\langle c(t)c(t+\tau)\rangle}{\langle c^2(t)\rangle} \quad \text{(S2a)}$$

$$= \frac{\langle \cos[2kz(t)]\cos[2kz(t+\tau)]\rangle}{\langle \cos^2[2kz(t)]\rangle} \quad \text{(S2b)}$$

$$= \frac{\langle \cos[2kz(t)]\cos[2k(\mathrm{z}+\Delta\mathrm{z}(\tau)]\rangle}{\langle \cos^2[2kz(t)]\rangle} \quad \text{(S2c)}$$

$$= \langle cos(2k\Delta z(\tau))\rangle \quad \text{(S2d)}$$

Where the final line arises from simplifying by trigonometric identities, including noting that ⟨sin4kz⟩ = 0. To evaluate the ACF, we consider the displacement probability distribution for 1-dimensional Brownian motion:

$$P(\Delta z|\tau) \sim exp\left(-\frac{\Delta z^2}{4D\tau}\right) \quad \text{(S3)}$$

To simplify evaluation, we introduce dimensionless parameters:

$$\widetilde{\Delta z} = 2k\Delta z(\tau) \quad \text{(S4)}$$

$$\tilde{\sigma}^2 = 8k^2 D\tau \quad \text{(S5)}$$

$$P(\widetilde{\Delta z}|\tau) = exp\left(-\frac{\widetilde{\Delta z}^2}{2\tilde{\sigma}^2}\right) \quad \text{(S6)}$$

This leaves us to evaluate ACF(t) = $\langle cos(Z)\rangle$ Where Z is a random variable drawn from $P(\widetilde{\Delta z}|\tau)$:

Taylor series expansion:

$$\langle cos(Z)\rangle = \left\langle \sum_{k=0}^{\infty}(-1)^k \frac{1}{(2k)!}(Z)^{2k}\right\rangle \quad \text{(S7a)}$$

Commuting sums:

$$= \sum_{k=0}^{\infty}(-1)^k \frac{1}{(2k)!}\langle (Z)^{2k}\rangle \quad \text{(S7b)}$$

Even moments of normal distribution:
$\langle (Z)^{2k}\rangle = \tilde{\sigma}(2k\text{-}1)\,\langle (Z)^{2k-2}\rangle$
$= \tilde{\sigma}^{2k}(2k-1)!!$
$= \tilde{\sigma}^{2k}(2k-1)(2k-3)\ldots 1$

$$= \sum_{k=0}^{\infty}(-1)^k \frac{1}{(2k)!}\tilde{\sigma}^{2k}(2k-1)!! \quad \text{(S7c)}$$

$$\frac{(2k-1)!!}{(2k)!} = \frac{(2k-1)(2k-3)\dots 1}{(2k)(2k-1)\dots 1} = \frac{1}{2^k k!} \qquad = \sum_{k=0}^{\infty} (-1)^k \frac{1}{2^k k!} \tilde{\sigma}^{2k} \tag{S7d}$$

$$= \sum_{k=0}^{\infty} \frac{\left(-\frac{\tilde{\sigma}^2}{2}\right)^k}{k!} = exp\left(-\frac{\tilde{\sigma}^2}{2}\right) \tag{S7e}$$

$$= \boldsymbol{exp\ (-4k^2 D\tau)} \tag{S7f}$$

### Note S2: ISABEL simulation details

Confinement effects on diffusion

The confinement effects on diffusion arise due to symmetry breaking near a wall and the attenuation of the long-range collective forces from the solvent acting on the particle. The relative diffusion attenuation is well-approximated in terms of the ratio *a/z* of the particle radius *a* and distance *z* from a wall:[1]

$$\lambda_{\perp} = \frac{6-10\left(\frac{a}{z}\right)+4\left(\frac{a}{z}\right)^2}{6-3\left(\frac{a}{z}\right)-\left(\frac{a}{z}\right)^2} \tag{S8}$$

$$\lambda_{\parallel}^{-1} = 1 - \frac{8}{15} ln\left(1-\frac{a}{z}\right) + 0.029\left(\frac{a}{z}\right) + 0.04973\left(\frac{a}{z}\right)^2 - 0.1249\left(\frac{a}{z}\right)^3 \tag{S9}$$

where $\lambda_{\perp}$ corresponds to the diffusion attenuation perpendicular to the wall (i.e. in *z*), and $\lambda_{\parallel}$ corresponds to the direction parallel to the wall (*x* and *y*).

To account for the attenuations from two walls ($\lambda_{w1}$ and $\lambda_{w2}$) separated by a height *h*, we apply the effect of each independently, known as the linear superposition approximation:[2]

$$\lambda_{2w}(z) = \lambda_{w1}(z) + \lambda_{w2}(h-z) - 1 \tag{S10}$$

Therefore for the ISABEL geometry, $D_z(z) = \lambda_{\perp,2w}(z)D_0$ and $D_{xy}(z) = \lambda_{\parallel,2w}(z)D_0$. The *z*-dependent $D_z$ and $D_{xy}$ values for a 100 nm bead in a 690 nm channel are shown below in Fig. S6, and the ACF accounting for confined diffusion of a model 100 nm particle is shown in Fig. S2.

### Optical forces

In addition to confinement, we incorporated the effect of optical forces on the particle. For each simulated particle, we could attribute a scattering and absorption cross sections $\sigma_{scat}$ and $\sigma_{abs}$ at 808 nm as determined by our past measurements.[3] For example, the 100 nm polystyrene beads exhibited an average $\sigma_{scat}$ = 36 nm$^2$ and $\sigma_{abs}$ = 0 nm$^2$, whereas a 50 nm Au bead exhibits $\sigma_{scat}$ = 52 nm$^2$ and $\sigma_{abs}$ = 52 nm$^2$ on average. To account for the range of $\sigma_{scat}$ values over a range of particle diameters $d_i$, we approximate a scaled scattering cross-section $\sigma_{scat,i}$ for an individual particle:

$$\sigma_{scat,i} = \sigma_{scat,mean}\,(d_i/d_{mean})^6 \tag{S11}$$

in accordance with the Rayleigh regime for scaling of scattering cross section. Mean scattering cross sections for each class of particles are listed in Table S2. Additionally, our scattering illumination beam was determined to have a power of 20 mW for most samples (7 mW for 50 nm Au NPs and 130 nm polystyrene NPs) and a 1/e$^2$ radius of 790 nm, leading to a peak intensity $I_{pk}$ 510 kW/cm$^2$ and average power 16 kW/cm$^2$ over the Knight's Tour pattern. From these quantities, we could calculate the scattering force acting on our particle:[4]

$$\boldsymbol{F}_{scat}(r) = \frac{n\sigma_{ext}}{c}\boldsymbol{S}(r) = \frac{n\sigma_{ext}}{c}I(r)\hat{\boldsymbol{z}} \tag{S12}$$

Where $\sigma_{ext} = \sigma_{scat} + \sigma_{abs}$ is the extinction cross section of the particle, $n$ is the refractive index of the medium (Table S1), and **S** is the Poynting vector of the beam. For a 100 nm PS bead, $\mathbf{F}_{scat}$ = 1.62 fN at the center of the beam. In the simulation, $|\mathbf{F}_{scat}|$ acting on the particle was calculated at each time step and used to calculate the $z$ placement of the particle as part of the third terms in Eq. 6-8 in the main text. Gradient forces were found to have negligible influence on the particle's contrast ACF. However, the scattering force was low magnitude but consistent, resulting in a bias where particles preferentially explored higher regions nearer the top wall. By biasing particles closer to the top wall, they experience an even slower $\langle D_z \rangle$ than expected due to confinement alone (Fig. S2).

### Interferometric scattering contrast model

To ensure efficient determination of the interferometric scattering contrast at each point, we implemented a contrast lookup table, which determined normalized interferometric scattering contrast $c(z, r)$ as a function of particle height $z$ and distance $r$ from the beam center $r_0$.

The model is based on the scattering of circularly polarized light from a dipole above a glass surface.[5-7] Consider a Rayleigh scatterer illuminated by a Gaussian beam described by the equation:

$$E_i(r,z) = \frac{w_0}{w_z} e^{-\frac{(r-r_0)^2}{w_z^2}} \exp\left(-ik(z-z_0) + k(z-z_0)^2 R_z^{-1} - \tan^{-1}\left(\frac{z-z_0}{z_r}\right)\right) \tag{S13}$$

Where $r = (x^2 + y^2)^{1/2}$ is the distance from the beam's optical axis at $r_0$, $w_0$ is the focal waist, $w_z$ is the waist a distance $z$ above the focus, and $z_0$ is the focal height of the beam with respect to the surface, which we took to be the midpoint between the two walls at $z$ = 345 nm. The $z$-dependent radius of curvature $R_z$ and Guoy phase appear in the penultimate and final terms in parentheses, respectively.

The reflected Gaussian beam is given by

$$E_r(r,z) = -r_q \frac{w_0}{w_z} e^{-\frac{(r-r_0)^2}{w_z^2}} \exp\left(-ik(z-z_0) + k(z-z_0)^2 R_z^{-1} - \tan^{-1}\left(\frac{z-z_0}{z_r}\right)\right) \quad \text{(S14)}$$

Where $r_q$ is the depolarized Fresnel reflectivity coefficient at the quartz-water interface. The negative sign prefactor indicates the reversal of propagation direction of the reflected beam. We can treat the reflection as from one surface (even though we have two surfaces in the trap) since it can be shown that the effect of two flat surfaces is a net reflected field that is merely a scaled version of the first reflection. As well, since the two surfaces are within the Rayleigh length of ~ 2μm, there is no significant defocus between the two reflections of the incident Gaussian beam.

Upon illumination, the Rayleigh scatter emits an electric field according to a dipole pattern with an orientation $\hat{\boldsymbol{\mu}}$ of the emitter dipole induced by the incident field. The emission from this induced dipole can then be modeled with a vectorial imaging model.[6,7] For a point-like particle illuminated by circularly polarized light, the dipole emission pattern appears radially polarized in the transverse direction, such that we can describe it as

$$\hat{\boldsymbol{\mu}} = \begin{bmatrix} \mu_x \\ \mu_y \\ \mu_z \end{bmatrix} = \frac{|\mu|}{\sqrt{2}} \begin{bmatrix} 1 \\ i \\ 0 \end{bmatrix} \quad \text{(S15)}$$

At the sample, the *s*- and *p*-polarized components of the radiated electric field are:[6]

$$E_s(\theta_2, \phi) = t_s \frac{\mathrm{n_1} \cos(\theta_1)}{\mathrm{n_2} \cos(\theta_2)} \left[\mu_y \cos(\phi) - \mu_x \sin(\phi)\right] \quad \text{(S16)}$$

$$E_p(\theta_2, \phi) = t_p \frac{\mathrm{n_1}}{\mathrm{n_2}} \left[\cos(\theta_1)\left(\mu_x \cos(\phi) + \mu_y \sin(\phi)\right) - \mu_z \sin(\theta_1) \frac{\mathrm{n_1} \cos(\theta_1)}{\mathrm{n_2} \cos(\theta_2)}\right] \quad \text{(S17)}$$

where $\theta_1$ and $\theta_2$ are polar angles of the emission in quartz and water, respectively and related by Snell's Law, $\phi$ is the emission's polar angle, $n_1$ and $n_2$ are the refractive indices in quartz and water, respectively, and $t_s$ and $t_p$ are the Fresnel transmission coefficients for *s* and *p* polarized light. A diagram of the coordinate system is provided in Fig. S11a.

These fields propagate through the objective to result in back-focal plane field components

$$E_{bfp,x}(\theta_1, \phi) = \frac{1}{\sqrt{\cos(\theta_1)}} \left[E_p \cos(\phi) - E_s \sin(\phi)\right] \exp[ik_1 z_0 \cos(\theta_1)] \exp\left[ik_2 z cos(\theta_2) + \frac{\pi}{2}\right] \quad \text{(S18)}$$

$$E_{bfp,y}(\theta_1, \phi) = \frac{1}{\sqrt{\cos(\theta_1)}} \left[E_s \cos(\phi) + E_p \sin(\phi)\right] \exp[ik_1 z_0 \cos(\theta_1)] \exp\left[ik_2 z cos(\theta_2) + \frac{\pi}{2}\right] \quad \text{(S19)}$$

Where the first bracketed term corresponds to the coordinate rotation to convert *s*- and *p*-polarizations to *x* and *y*, the first exponential corresponds to the phase delay associated with the beam's focal height, and the second exponential corresponds to the phase delay due to the height of the particle and π/2 phase delay due to nonresonant scattering. The back-focal-plane field is apertured according to the NA of the objective. A representative back-focal plane intensity $|E_{bfp,x} + E_{bfp,y}|^2$ for a scatterer at $z$ = 350 nm is shown in Fig. S11b.

Finally, the dipole emission pattern in the image plane conjugate to the sample plane is the 2D Fourier transform of the back-focal-plane field:

$$\boldsymbol{E}_{\text{scat}}(x,y) = FT\left\{\begin{bmatrix} E_{bfp,x}(\theta_1,\phi) \\ E_{bfp,y}(\theta_1,\phi) \end{bmatrix}\right\} \tag{S20}$$

The scattering intensity $|E_{\text{scat,x}} + E_{\text{scat,y}}|^2$ in the sample plane is shown in Fig. S11c.

With the scattering and reflected fields determined, the interferometric scattering contrast was calculated according to:

$$c = \frac{\sum_{x,y}|E_r(x,y)+E_{scat}(x,y)|^2 - \sum_{x,y}|E_r(x,y)|^2}{\sum_{x,y}|E_r(x,y)|^2} \tag{S21}$$

where the summations arise from detection on a single-element photoreceiver.

Finally, to assemble the contrast map, we repeated the above Gaussian beam and dipole scattering calculations over a grid of $z \in$ [0: 10: 690] nm and $r \in$ [0:40:800] nm, further interpolated to a grid spacing of 3.45 nm in $z$ and 8 nm in $r$. The contrast map was then normalized to the value of highest absolute contrast. During a Brownian Dynamics simulation, the particle's $z$ position and distance $r$ from the scanning beam was determined at each time step and matched to the corresponding position to extract its contrast $c(t)$.

#### Note S3: Characterization of ABEL Cell Depth

To accurately model the hydrodynamic confinement effects felt by each nanoparticle, we required an accurate estimation of the microfluidic cell height. We achieved this by white light interferometry, where the reflected spectrum of a white light source (Leukos Rock 400 supercontinuum) displayed oscillations due to interference between the two surfaces. The illumination geometry is shown in Fig. S3 below.

We measured the reflected spectrum from the trapping region of an ABEL cell, normalized by the reflection of a single quartz surface to retrieve the fringes as a function of wavelength. The interference of two reflections $I_{\text{r,tot}}$ can be described by the following equation:

$$I_{r,tot}(\tilde{\nu}) = \cos(4\pi n\,\tilde{\nu} h) \tag{S22}$$

Where $\tilde{\nu}$ is the spectroscopic wavenumber, $n$ is the index of refraction in the gap, and $h$ is the distance between the two walls. We measured the reflected spectrum with an air gap such that $n$=1. As shown in Fig. S4, the ABEL cell used for the bead standards exhibited $h$ = 690 nm gap, whereas the cell used to trap carboxysomes showed a height $h$ = 1260 nm.

**Table S1: Universal simulation parameters**

| Parameter | Value |
|---|---|
| Temperature *T* | 19.0 °C |
| Viscosity *η* | 1.03 mPa·s |
| Beam Wavelength *λ* | 808 nm |
| Medium refractive index *n* | 1.33 |
| Beam Waist $w_0$ | 790 nm |
| Beam axial focal position $z_f$ | 345 nm |
| Feedback gain *g* | 2.28 V/μm |

**Table S2: Particle-specific simulation parameters**

| Particle | Electrokinetic Mobility *μ* (μm $V^{-1}$ $s^{-1}$) | Mean scattering cross-section $\sigma_{scat}$ ($nm^2$) | Laser Power *P* (mW) | Time Step Δ*t* (μs) | Microfluidic Height *h* (nm) |
|---|---|---|---|---|---|
| 100 nm polystyrene | 20 | 36 | 20 | 3.75 | 690 |
| 130 nm polystyrene | 20 | 120 | 7 | 3.75 | 690 |
| 80 nm polystyrene | 30 | 7 | 20 | 3.75 | 690 |
| 50 nm gold | 40 | 52 | 7 | 3.75 | 690 |
| Carboxysomes | 20 | 2 | 20 | 18.75 | 1260 |

### Supplementary Figures

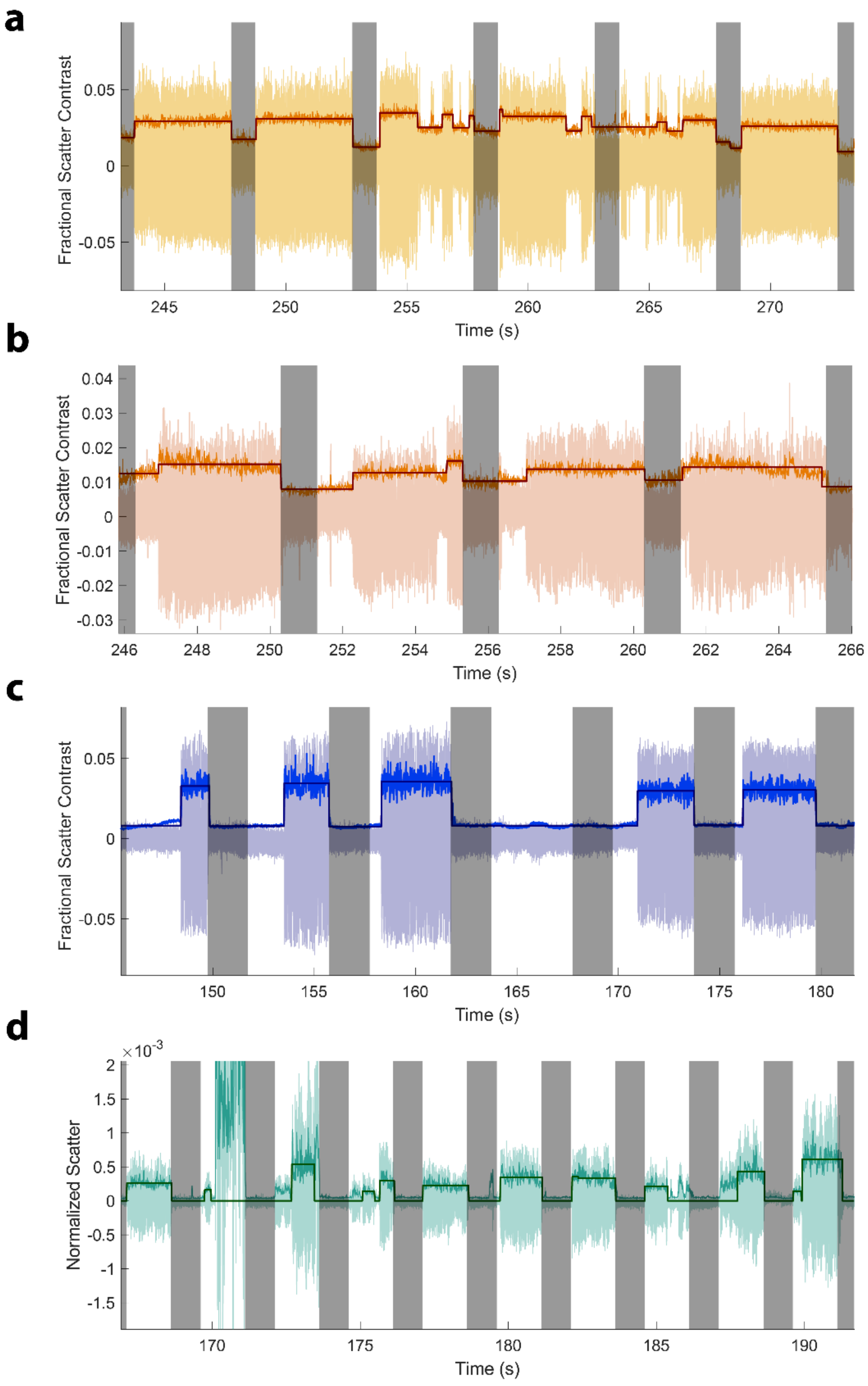

**Figure S1:** Sample trapping trajectories from additional nanoparticles: (a) 50 nm Au beads, (b) 80 nm polystyrene beads, (c) 130 nm polystyrene beads, and (d) carboxysomes.

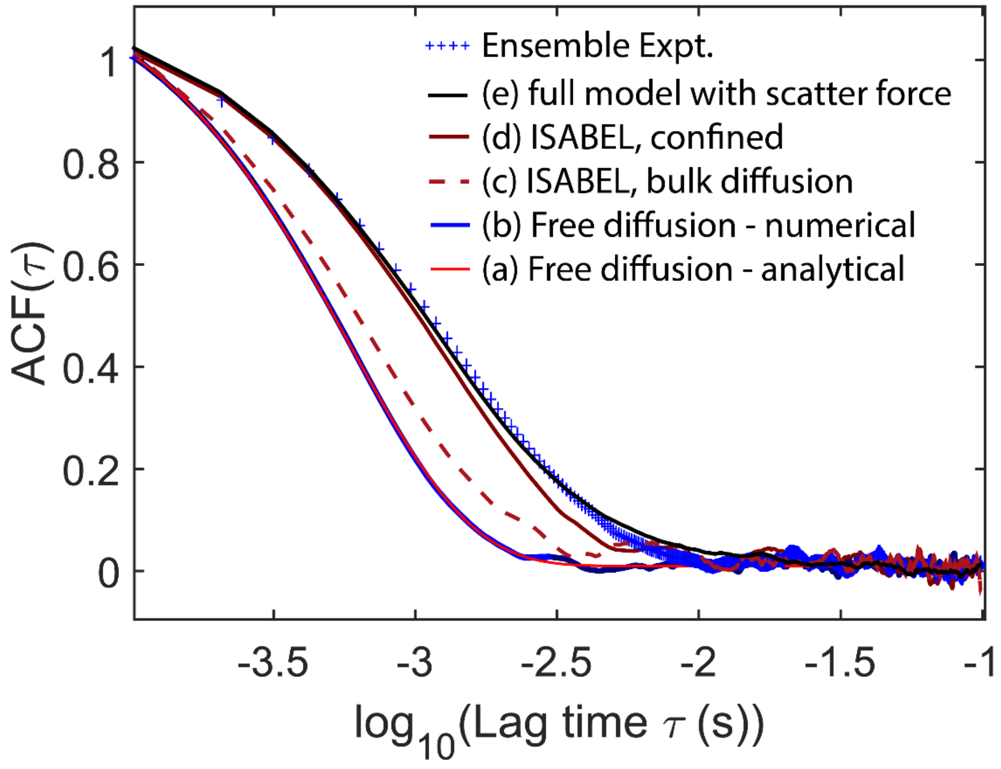

**Figure S2:** ACF curves for free diffusion of a 110 nm bead versus contributions to the ISABEL experiment. (a) Analytical free diffusion in one dimension (b) numerical model, (c) contrast ACF in the ISABEL trap for a particle subject to an isotropic diffusion coefficient (no confinement), (d) ISABEL trap with confinement effects, and (e) full model, including optical scattering forces.

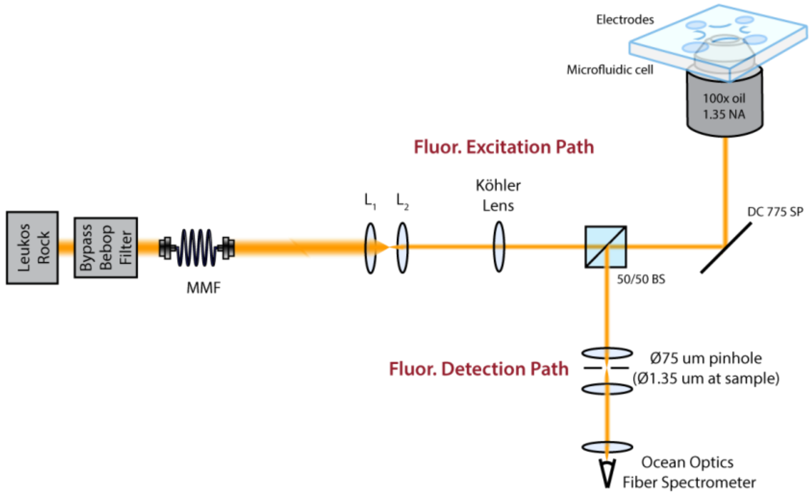

**Figure S3:** White-light illumination path to determine the ABEL microfluidic height *h*.

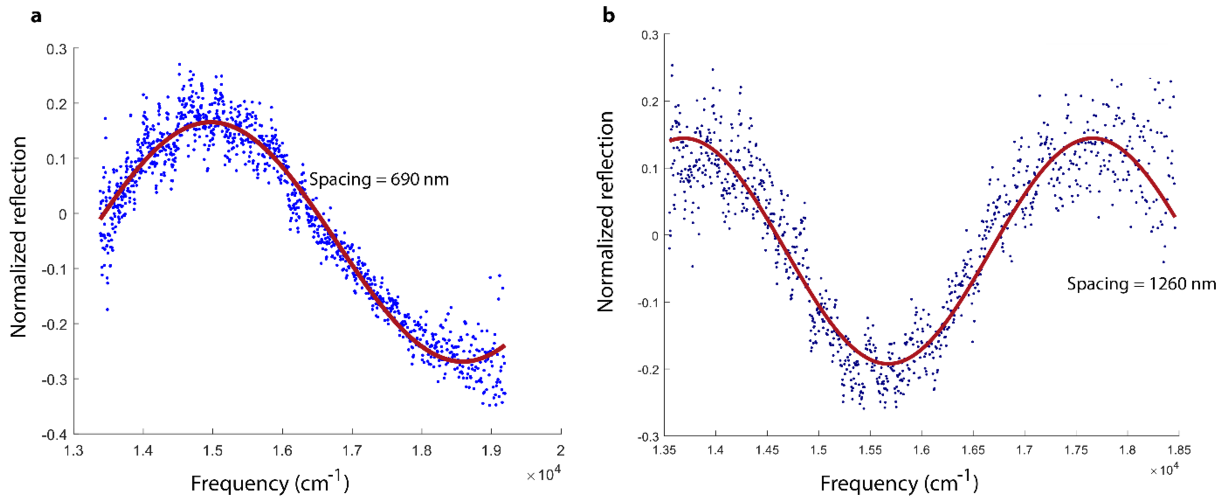

**Figure S4:** Normalized reflected interference and sinusoidal fit as a function of optical frequency. (a) Reflectivity in the microfluidic cell used for bead standards. (b) Reflectivity for the microfluidic cell used to trap carboxysomes.

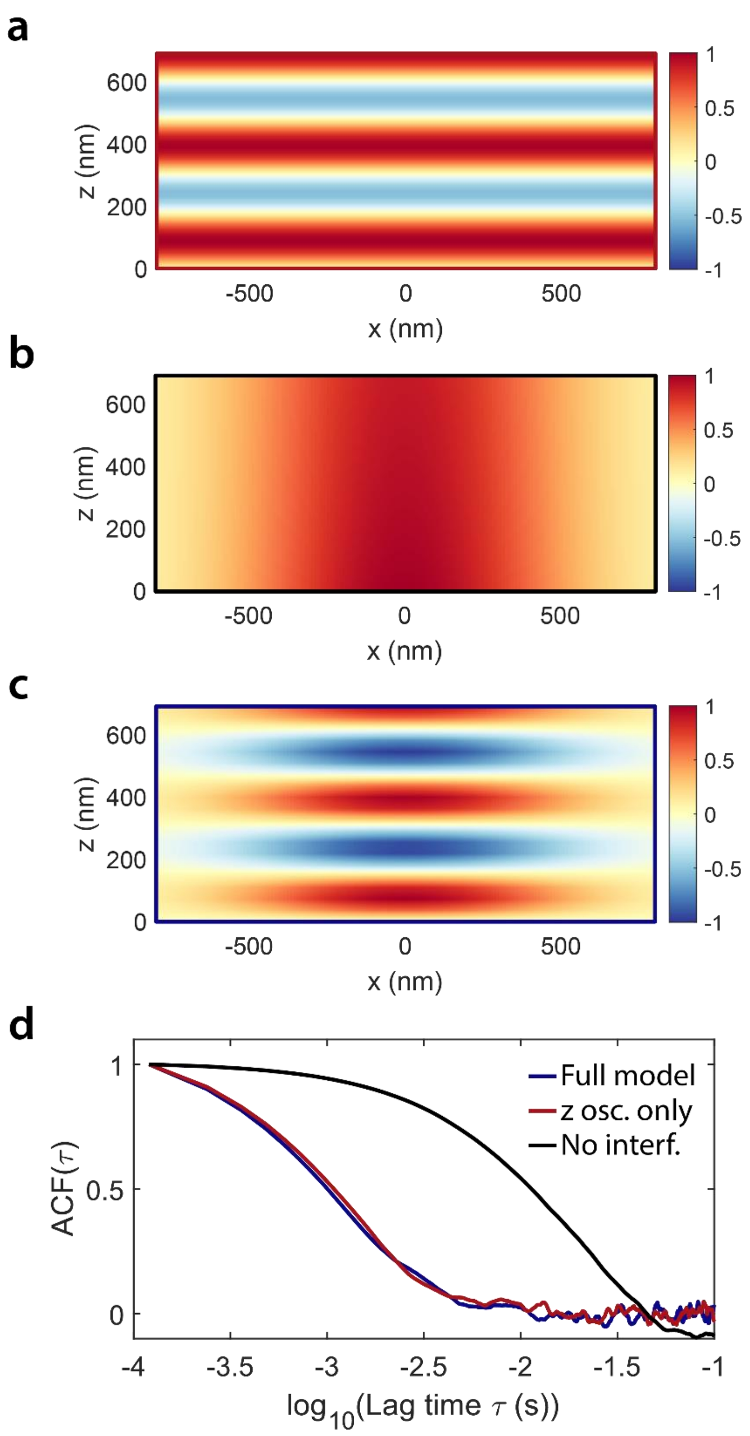

**Figure S5:** Assessment of the lateral vs axial diffusion components in the contrast ACF. (a) Contrast map with only interference, generated by removing the Gaussian beam waist contributions in Eq. S13 and S14. (b) Contrast map without interference effects. (c) Full contrast model. (d) Simulated ACFs for a trapped 110 nm bead under each contrast model.

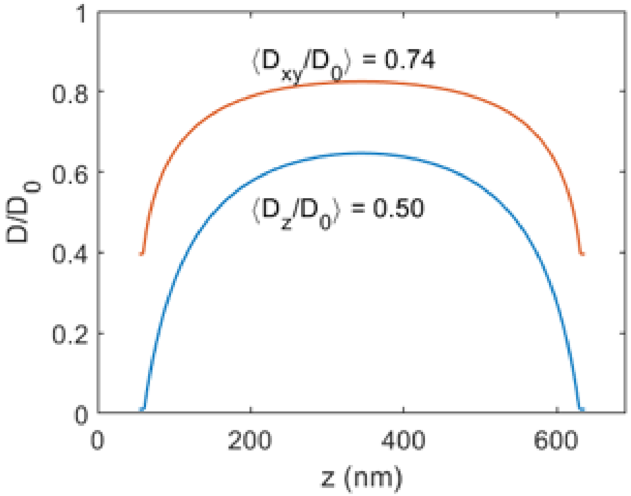

**Figure S6:** The diffusion attenuation experienced by a $d_H$ = 100 nm particle between two walls separated a distance $h$ = 690 nm. The orange curve corresponds to lateral diffusion with the average attenuation shown in brackets, while the blue curve shows axial diffusion. Diffusion attenuation is strongest near the walls, and the axial motion is more heavily attenuated by the presence of two walls.

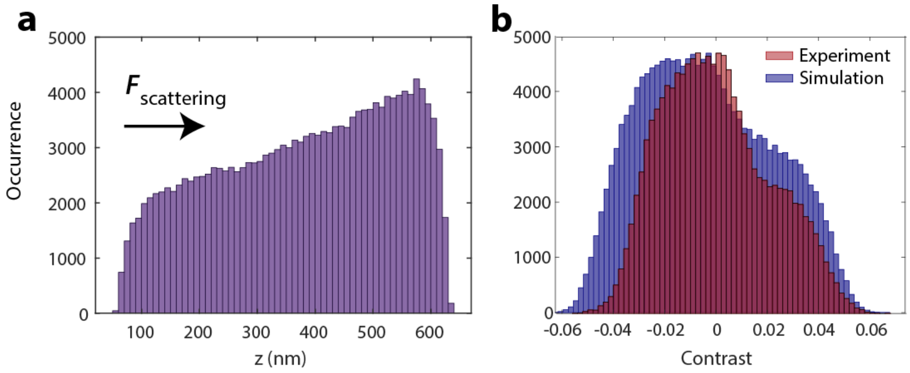

**Figure S7:** Influence on optical scattering on (a) distribution of $z$ positions of a 110nm polystyrene particle and (b) on asymmetry of the contrast histogram.

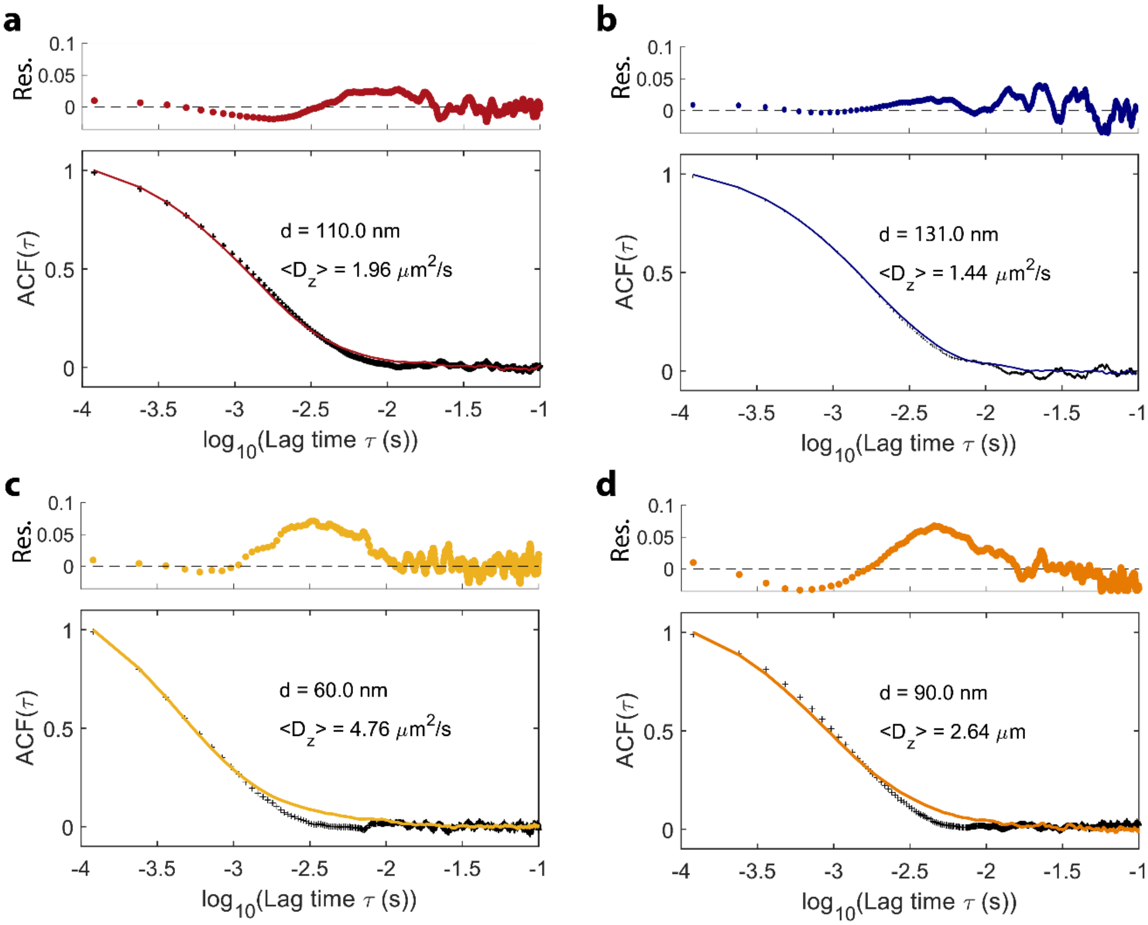

**Figure S8:** Comparison between numerical simulations of standard bead contrast ACFs with 10-bead ensemble-averaged measured ACFs: (a) 100 nm polystyrene beads, (b) 130 nm polystyrene beads, (c) 50 nm gold beads, and (d) 80-nm polystyrene beads. We note that the curves are not fits to experimental data, but generated from simulations with parameters listed in Tables S1 and S2.

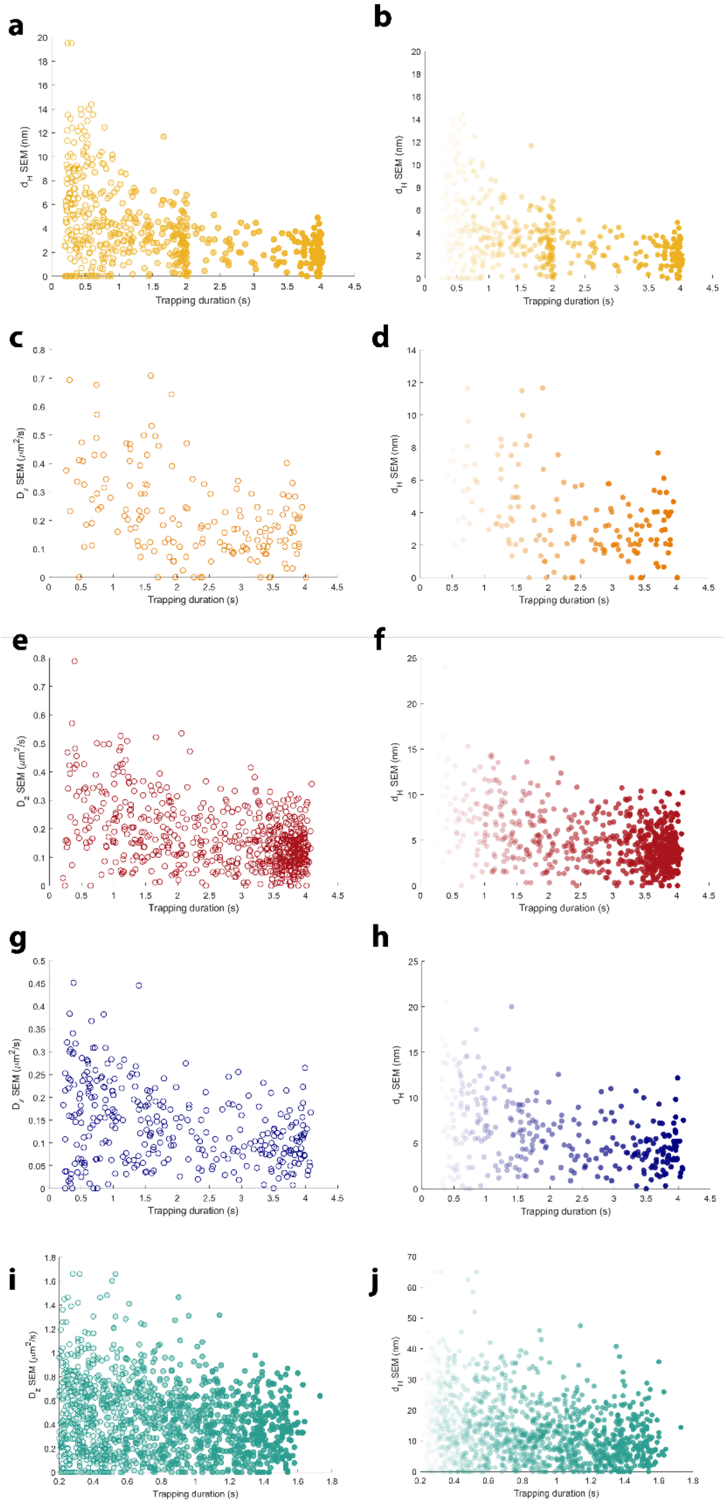

**Figure S9:** Standard errors of the mean (SEMs) for ⟨$D_z$⟩ (left column) and $d_H$ (right column) estimates for all particles trapped. (a)-(b) 50 nm Au beads, (c)-(d) 80 nm polystyrene, (e)-(f) 100 nm polystyrene, (g)-(h) 130 nm beads, and (i)-(j) carboxysomes.

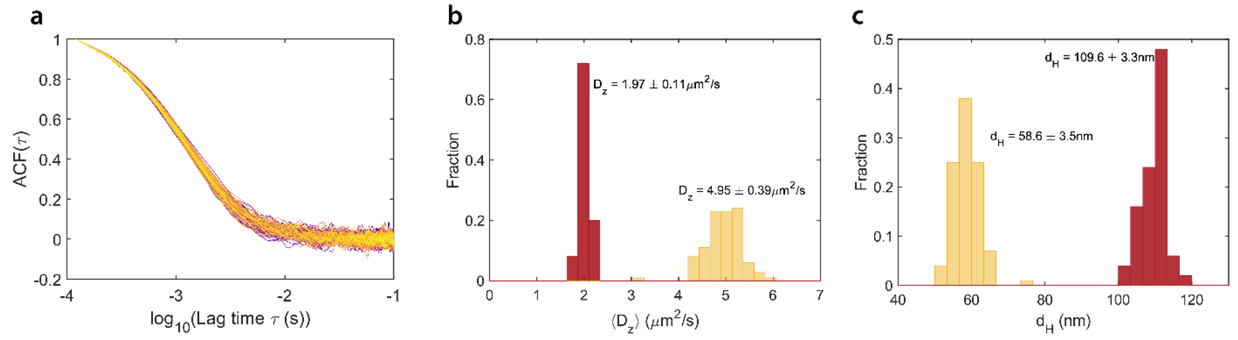

**Figure S10:** Benchmarking ACF fitting precision and accuracy with 100 identical model particles of fixed $d_H$ for 110 nm polystyrene particles and 59 nm Au particles. Simulations were conducted for 4s trapping events and recorded without experimental noise. (a) ACFs from 110nm beads, demonstrating variation in the ACF decay time due to finite statistical sampling over the trapping interval. (b) $D_z$ results and (c) $d_H$ results.

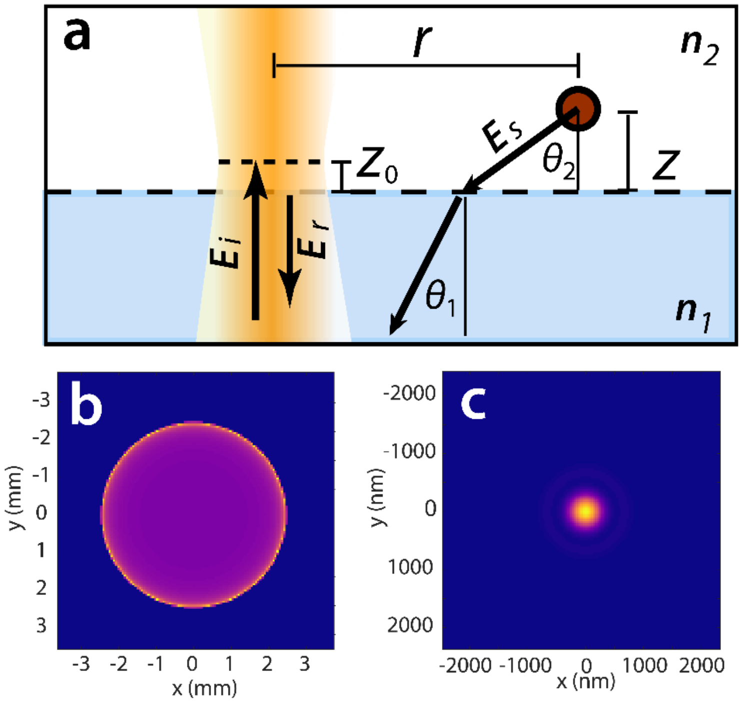

**Figure S11:** (a) Coordinate system and sample electric field calculations in the (b) back focal plane (magnification 100x) and (c) effective image plane used for the contrast model, plotting the intensity of the scatterer PSF. In (a), the polar angle $\phi$ of the dipole emission is not shown, and the beam-particle distance $r$ is not to scale.

### Supplementary References

1 X. Bian, C. Kim and G. E. Karniadakis, *Soft Matter,* 2016, **12**, 6331–6346.

2 B. Lin, J. Yu and S. A. Rice, *Phys. Rev. E,* 2000, **62**, 3909–3919.

3 A. A. Lavania, W. B. Carpenter, L. M. Oltrogge, D. Perez, J. B. Turnšek, D. F. Savage and W. E. Moerner, *J. Phys. Chem. B,* 2022, **126**, 8747–8759.

4 G. Pesce, P. H. Jones, O. M. Maragò and G. Volpe, *Eur. Phys. J. Plus,* 2020, **135**, 949.

5 M. A. Lieb, J. M. Zavislan and L. Novotny, *J. Opt. Soc. Am. B,* 2004, **21**, 1210–1215.

6 A. S. Backer and W. E. Moerner, *J. Phys. Chem. B,* 2014, **118**, 8313–8329.

7 D. Axelrod, *J. Microsc.,* 2012, **247**, 147–160.
